# Charting the molecular landscape of neuronal organisation within the hippocampus using cryo electron tomography

**DOI:** 10.1101/2024.10.14.617844

**Authors:** Calina Glynn, Jake L.R. Smith, Matthew Case, Rebecca Csöndör, Ana Katsini, Maria E. Sanita, Thomas S. Glen, Avery Pennington, Michael Grange

## Abstract

Cellular cryo-electron tomography (cryoET) enables the capture of detailed structural information within a biologically relevant environment. However, information in more complex samples, such as tissues, is lacking. Importantly, these observations need to be set in context of populations; imaging on the molecular scale to-date is limited to few observations *in-situ* that struggle to be generalised. This is due to limitations in throughput and versatility employed by current instrumentation. Here, we utilise plasma focused ion beam milling to examine the molecular landscape of mouse hippocampus by cryoET in targeted regions across multiple individuals revealing the complex organisation of macromolecules from the CA1 strata pyramidale (sp) to radiatum (sr). Our data represent a molecular atlas, producing snapshots of hippocampal architecture in adult mouse. The combination of instrumentation and application of technical advancements provides a framework to explore specific structural questions within native tissues in a targeted manner.

## Introduction

Within the brain, a myriad of molecular interactions are responsible for maintaining cellular connectivity, homeostasis and maintenance. Characterising the molecular and cellular organisation of molecules within the context of brain tissue could unlock fundamental mechanistic insight into a range of processes, with consequences for our understanding of health and disease. Structures of proteins, complexes, and other macromolecules have enabled us to understand the molecular underpinning of many processes. Very few approaches exist that enable a molecular assessment of macromolecules in the native context of an intact brain, including within biopsies. Routine tissue analysis remains challenging, particularly when investigating molecular aspects across multiple subjects to understand disease-related genotype-phenotype relationships. Tools that bridge the gap between structure, function and cellular dysfunction are needed.

To investigate molecules in context, the vitrification state is crucial as biological molecules require hydration to remain in a folded, functional state. Provided this requirement is met, it is now possible to visualise cellular architecture and determine the structures of macromolecules to sub-nanometre resolution by sub-volume averaging^1–3^. Tissues present a particular challenge as biopsy section thickness is ∼1000 fold greater than the 200-300 nm sectioning required to create electron transparent regions suitable for imaging on the molecular scale via cryo-electron tomography (cryoET). Cryo-ultramicrotomy (CEMOVIS) can be used to section ribbons of tissue down to 40 nm in thickness^4^ but suffers from multiple artefacts^5–8^ obfuscating detail. Cryo-focused ion beam (FIB) milling is now routinely used for creating thin lamellae of cellular samples^9–12^ however conventional traditional liquid metal ion source (LMIS) FIBs are limited in sputter yield but have been demonstrated to work with cryo lift-out of samples ∼ 50 µm depth^13^ Therefore, for tissue samples, where thickness is typically ≥ 100 µm, larger volumes of material need to be removed to fabricate lamellae. Alternatively, plasma focused ion beams (PFIB) have recently been utilised for cryogenic life science samples^14^, both for volume electron microscopy^15^ and lamellae fabrication, allowing for high-resolution sub-tomogram averaging^16,17^. The increased sputter yield of xenon plasma allows for excavation of large volumes, necessary for lamellae fabrication in thick specimens in a shorter time frame^16,18^.

Lamellae fabrication from samples high-pressure frozen directly onto electron microscopy grids has been demonstrated^19,20^. For tissues, however, this results in compression of material during high pressure freezing (HPF), deforming morphological features within the native tissue^20^. Alternatively, cryo-lift-out has been demonstrated to generate spatially related lamellae across entire organisms and multicellular samples^21^ though due to the use of LMIS, this is still limited to depths of sample 30–50 µm thick. As methods to section tissues can cause physical damage to cells near the surface of the tissue section, increasing the depth at which tissues can be excised for structural biology approaches will increase its effectiveness at investigating regions of interest on the molecular scale.

Here we demonstrate the ability to robustly vitrify mouse brain tissue within three hours post-mortem for structural characterisation by cryoET. Utilising the well-characterised global morphology of the mouse hippocampus to map each layer using cryo-fluorescence microscopy, we have adapted a PFIB milling and cryo lift-out strategy to image specific sub-regions of the CA1 region of the hippocampus from samples frozen in HPF carriers up to 200 µm thick. This enabled features, particularly synapses and the apical dendrite network, to be characterised within this layer by cryoET. We demonstrate this using biopsies from several mice and by sampling multiple regions within the brain, exemplifying the potential to investigate targeted structural biological questions across cohorts. We further demonstrate its versatility by incorporating different sample geometries, using a planar lift-out to generate serial sections and lamella spanning from the CA1 strata pyramidale through radiatum, maintaining spatial information on the molecular scale. This illustrates the capacity to apply high-throughput lamellae fabrication and cryo-ET analysis to characterise the molecular landscape within tissues. The basis of this work forms a framework for the future investigation of the molecular organisation crucial to disease, not just of the brain, across multiple individuals.

## Results

### Extraction of brain tissue from high pressure freezing carriers

One of the main challenges of producing tissue samples for cryoET imaging is to manipulate and thin vitrified biopsies (∼200 µm) to electron transparent sections (∼200 nm) in a timely manner, without altering their structure. We were able to develop two routines, “perpendicular” and “planar”, enabling different geometries for lift out (Figure 1a). This used an adapted cryo-lift-out^21^ approach to shape and extract regions preserving the hydrated state of the sample using HPF. Prior to HPF, mouse hemibrains (∼6 months of age) were sectioned to 100-200 µm in thickness using a vibratome (Extended Data Table 1). Subsequently, sub-regions of hippocampus could be targeted and correlated using micron-scale fluorescence microscopy to nanometre-scale imaging by cryo electron tomography (cryoET) (Figure 1).

**Figure 1.**
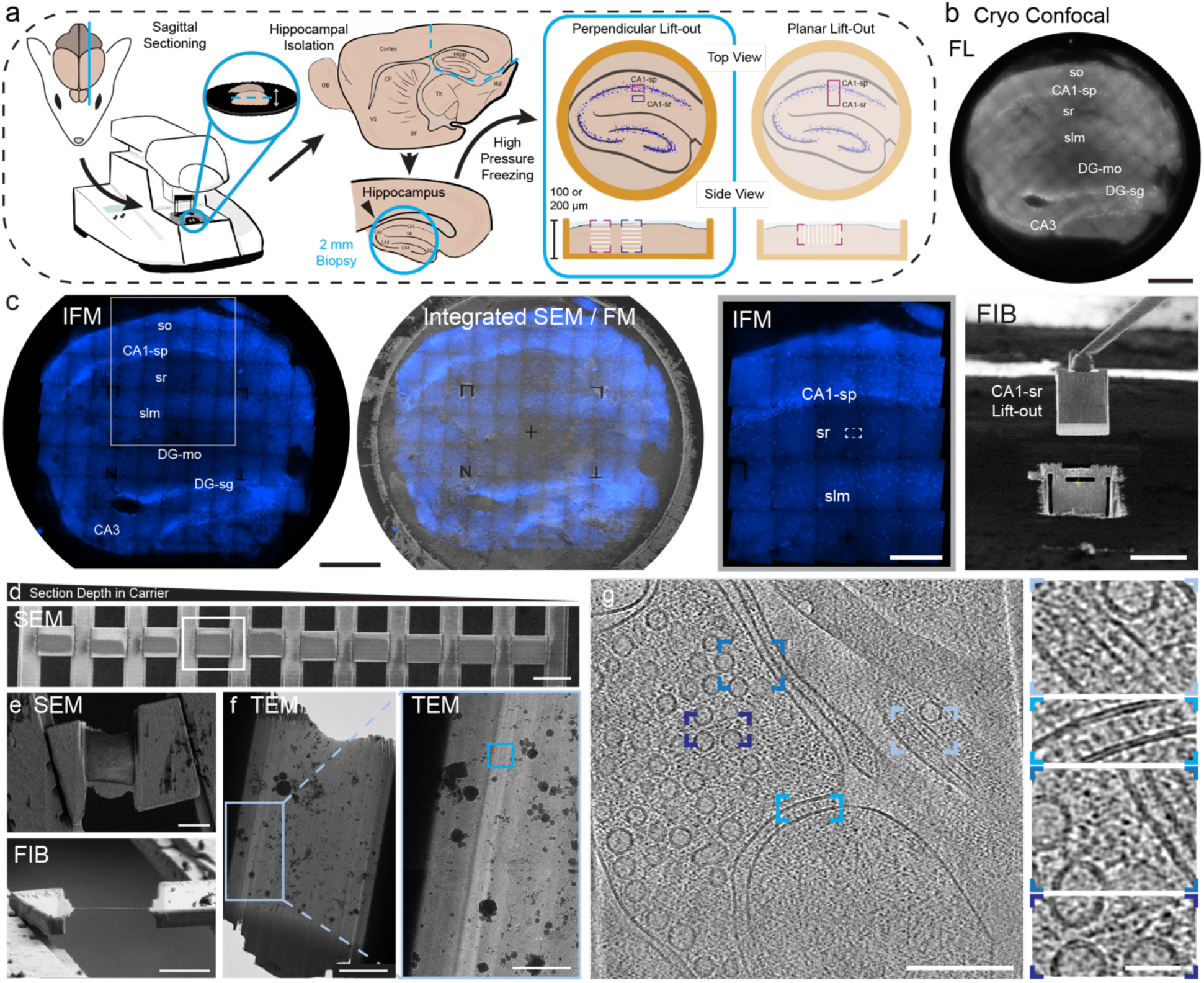
CLEM workflow for targeting features within vitrified mouse brain tissue. a) schematic of mouse brain sectioning and biopsy illustrating capacity to target specific layers within defined sections of mouse hippocampus for subsequent lift-out in perpendicular or planar geometries. In the remainder of this figure, a perpendicular lift-out (boxed) was used. b) Cryo-confocal image of a vitrified mouse hippocampal section in a high-pressure freezing carrier. Scale bar 500 µm. c) The same section imaged using the integrated fluorescence module (IFM) in the FIB/SEM (left, scale bar 500 µm) with CA1-sr lift-out target and lift-out marked (right, IFM scale bar 200 µm, FIB scale bar 50 µm). d) Serial sections from the CA1-sr lift-out with sections deeper in the carrier on the left and closer to the tissue surface at the right. Scale bar 50 µm. e) Thinned section boxed in (d) from SEM (top) and FIB (bottom) view. Scale bars 10 µm. f) Lamella of the same section in the TEM (scale bar 5 µm) with boxed region enlarged (right, scale bar 2 µm). g) Reconstructed tomogram from region boxed in the right image of f) showing a synapse (scale bar 200 nm) with details such as microtubules (top right), densities in the synaptic cleft (second from top, right), and densities spanning between synaptic vesicles and membranes (second from bottom and bottom right). Scale bar for insets in (f) 40 nm.

To excise regions of interest we used PFIB milling, which enables greater sputter yields. This is essential as extensive trench preparation and undercuts are required (Extended Data Figure 1). For a desired perpendicular lift length of ∼60-70 µm, 10-14 sections could be deposited with time taken for the various steps outlined in Extended Data Table 2. For planar geometries, the trench milling, and undercuts are larger due to the increased size and orientation, increasing the amount of time needed (Extended Data Table 2). For this work, 26 lift-outs were extracted resulting in 186 serial sections being deposited. Of these, 170 were retained (91.4%) upon transfer between microscopes for lamellae fabrication.

After deposition, serial sections were subsequently thinned to electron transparency using a combination of xenon plasma (to around ∼ 400 - 600 nm) and manually polishing with argon plasma. This hybrid approach was used as xenon has a milling rate 3-4 times greater than argon, reducing milling time for bulk removal of material before polishing with the finer argon beam^16^. Resulting lamella reached thicknesses comparable to those obtained for cellular lamella and had contrast transfer functions which could be accurately fit to sub nanometre resolution (Extended Data Figure 2).

### Brain tissue vitrification

To adequately carry out cryoET and structural studies in any environment, the sample needs to be maintained in an amorphous, frozen-hydrated state. Furthermore, the tissue needs to be maintained with minimal cell death and preserved without compression during high pressure freezing and in buffer conditions that lead to as little necrosis and morphological disturbance as possible. We could therefore investigate the optimal conditions for vitrification of mouse cortex and hippocampal sections.

We assessed the vitrification of mouse brain slices under different conditions, such as tissue thickness, cryoprotectant, and incubation conditions (Extended Data Figure 3, Extended Data Table 1). Vitrification was assessed based on the visibility of non-vitreous ice in transmission electron microscope (TEM) overview images of lamella, ice diffraction in TEM images, and distortion of membranes. We assessed 18 conditions through this approach and were able to determine several conditions where vitreous samples could be obtained. Importantly, several conditions led to partially vitrified preparations where the nature of the non-vitreous state could only be visualized through reflections in certain images (Extended Data Figure 3). Conditions that produced the most robust vitrification were incubated for at least 15 minutes with combinations of cryoprotectants of at least 10% low molecular weight (here, sucrose MW = 0.34 kDa or ethylene glycol, MW = 0.062 kDa) and higher molecular weight (e.g. dextran, MW = 40 kDa) components. Cryoprotectants containing only large MW components such as dextran or BSA (molecular weight 66.5 kDa) were insufficient to yield vitrified tissue, where samples exhibited either dehydration artefacts, such as distorted membranes, or contained reflections in tilt images at different angles (Extended Data Figure 3b). Tilt series for subsequent analysis were only acquired in lamellae where we did not observe any of these features.

Within vitrified sections of mouse cortex, we were able to observe common cellular and brain-specific features. These included myelin, synaptic vesicles, mitochondria, membranes, ribosomes, microtubules, and open space outside of membranes (Extended Data Figure 4). Interestingly, the substantial spaces between cells contrasts with chemically fixed tissues, where a large portion of the extracellular space is lost due to fixation^22^.

### Hippocampal layer targeting using cryo-CLEM

Using the known architecture of mouse hippocampus, we targeted the CA1 strata pyramidale (CA1-sp) and radiatum (CA1-sr) by mapping the location of large, neuronal, cell bodies with live-cell nuclear stain (Figure 1b, c). Cryo-confocal microscopy overviews (Figure 1b) enabled orientation information to be acquired prior to fluorescence mapping in the FIB/SEM chamber. Additionally, the slice integrity post-freezing and presence of a wide enough range of hippocampal features could be assessed for accurate downstream targeting. The orientation information was subsequently used for positioning the carrier for milling and correlation of fluorescence with the integrated fluorescence microscope (IFM) for targeted lift-out (Figure 1c). To map the cellular organisation of the entire tissue section, we acquired fluorescence tilesets that spanned the entire 2 mm by 2 mm carrier. With a 20x fluorescence objective, these tilesets could be acquired in as little as 20 minutes for one fluorophore depending on the desired number of Z steps (Extended Data Table 2).

We initially targeted, lifted-out, sectioned, and thinned regions of tissue containing fluorescently labelled cell bodies from the CA1-sp (Extended Data Figure 5), as this enabled us to validate our approach for fluorescent targets. Individual nuclei, including regions of heterochromatin and euchromatin, could be identified within the cryo-fluorescence data (Extended Data Figure 5a, c). Retention of fluorescent targets could be monitored at all stages, from initial carrier overviews and Z stacks through to the final thinned lamella to aid in targeted tilt series acquisition in the TEM. In this hippocampal layer, we were able to collect tilt series where most of the field of view was ribosomes along with tomograms containing mitochondria and vesicles, in line with features expected of and near cell bodies (Extended Data Figure 5).

Beyond the stratum pyramidale, we were also able to target sublayers that were largely devoid of fluorescence using the established morphology of mouse hippocampus. In this case, we selected the CA1-sr. This layer is known to be rich in synaptic connections^23^ and can be targeted via proximity to the CA1-sp, which was readily identified when the CA3 stratum pyramidale and the dentate gyrus were present in sections (Figure 1b). For targeting the CA1-sr, we first considered the positioning of the underlying dentate gyrus, the CA1 stratum lacunosum molecular (CA1-slm), and measured the thickness of the overlying CA1-sp nuclei (∼50 um). We targeted 50-150 µm below the CA1-sp nuclei for our CA1-sr lift-outs (Figure 1c).

From this sublayer, we collected six datasets originating from 2 mice, one male and one female, totalling 359 tilt series (Extended Data Table 3). In contrast to the CA1-sp datasets, we did not observe large numbers of ribosomes. As specific mechanisms exist to keep ribosomes localised to cell bodies, rather than migrating along neuronal processes, this is consistent with characteristics expected of the CA1-sr. Instead, we observed cellular features more typically observed in neuronal processes. These included mitochondria with (13/359, or 4%) and without (119/359, or 33%) granular deposits^24,25^, cytoskeletal elements like microtubules (302/359, or 84%) and thinner filaments including actin (114/359, or 32%) in dendrites and other processes, small clusters of 1-20 ribosomes (60/359, or 17%), processes with vesicles (311/359, or 87%), and synapses (56/359, between 3-30% of tomograms collected from each dataset) (Figure 1 and 2, Extended Data Figure 6 and 7, Extended Data Table 3). In our CA1-sr data we found abundant synapses where cell adhesion molecules and inter-membrane interactions could be seen (Figures 1g and 2).

**Figure 2.**
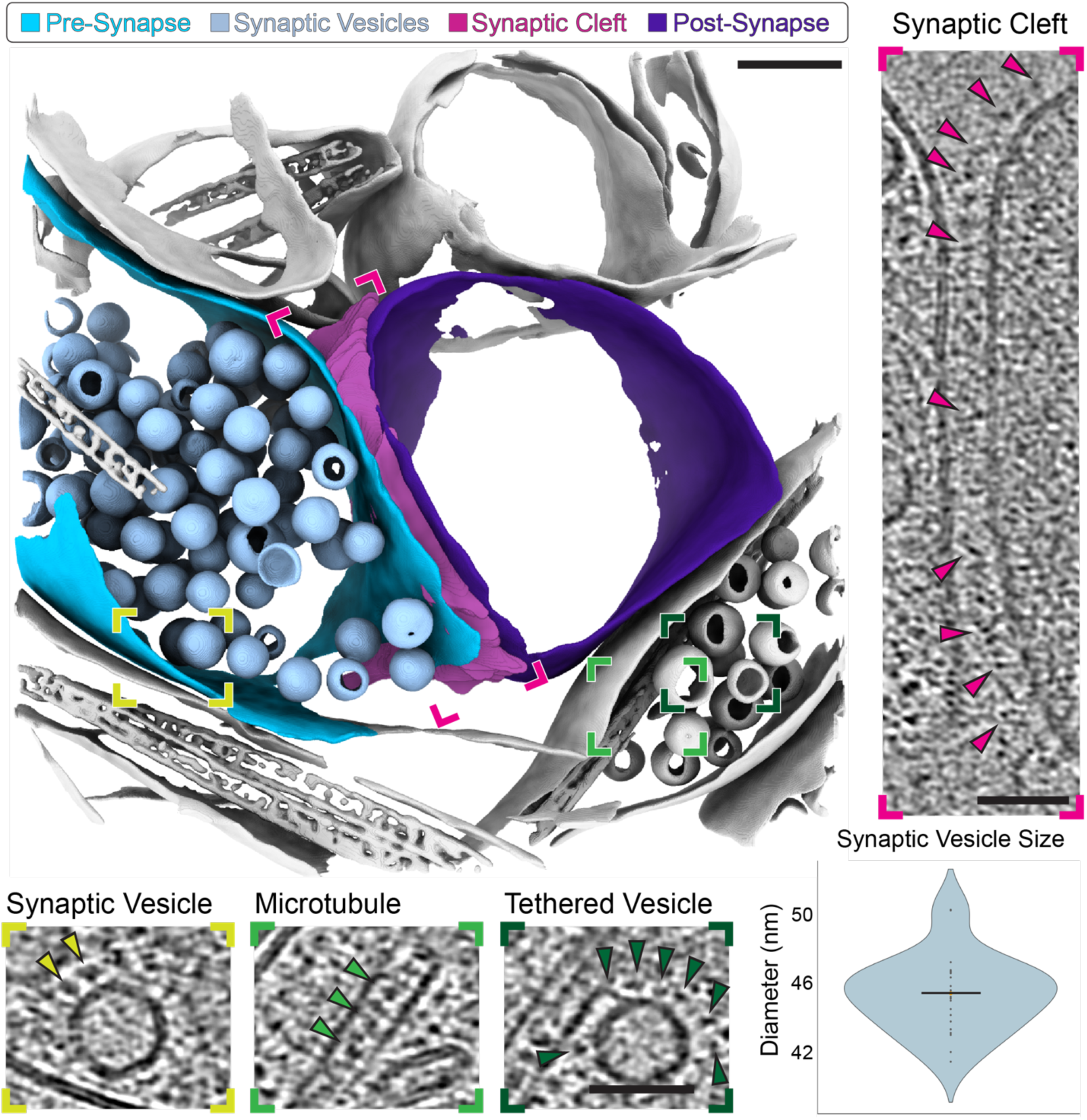
Synapse Segmentation. Segmentation of a synapse with synaptic vesicles (light blue), synaptic cleft (pink), pre-synaptic membrane (blue), and post-synaptic membrane (purple) highlighted. Scale bar 100 nm. Insets from the deconvolved tomogram highlighting details like synaptic cleft densities (right, pink arrows), synaptic vesicles with membrane attached densities (yellow, left and dark green, right arrows) and microtubules with interior densities (middle green arrows). Scale bar in insets (50 nm). Distribution of synaptic vesicle diameters across a subset of 27 tomograms where the only vesicles present are in a visible synapse (bottom right). Each dot represents the per-tomogram mean of the median of the per-slice 2d feret diameter of each vesicle.

### Synapse Organization within the CA1-sr

Synapses could be identified by their canonical organisation. This consisted of a presynapse, containing vesicles, and in some cases mitochondria, inside an enclosed membrane that widened ending with a cleft. This cleft contains multiple putative adhesion molecules spanning it. Finally, at the other side of the cleft is a postsynapse where the widened compartment narrows. Various subcellular organelles and molecular features could be visualised within the post-synaptic compartment. Of note, actin filaments and post-synaptic membrane densities could be observed (Extended Data Figure 7).

Within the presynapse, synaptic vesicles appeared spherical and could be further rendered in segmentations (Figure 2). Synaptic vesicles had a mean diameter of 45.4 ± 2.2 nm (Extended Data Figure 8), consistent with previously reported values for cryogenically preserved synaptic vesicles^26,27^ and larger than vesicles in resin embedded synapses^28^. Interestingly, we were able to observe the process of synaptic vesicle fusion with the presynaptic membrane in some instances of our data (Figure 3), evidencing synaptic activity occurring at the time of high pressure freezing. These events were somewhat rare and were noted in 8 out of 107 total synapses across all datasets. Indeed, the process of synaptic vesicle fusion is a fast process (∼ 100 µs^29^) with respect to high pressure freezing (∼ 50 ms). In the examples we acquired (Figure 3), the synaptic vesicle first forms tethers with the presynaptic membrane, often in proximity to apparent tethering densities that could be recruitment molecules such as Munc13-1^30^. Besides the presynaptic vesicles, we could also occasionally identify large dense core vesicles on the presynaptic side, which could be identified by their large size and electron dense interior^25^ (Extended Data Figure 7).

**Figure 3.**
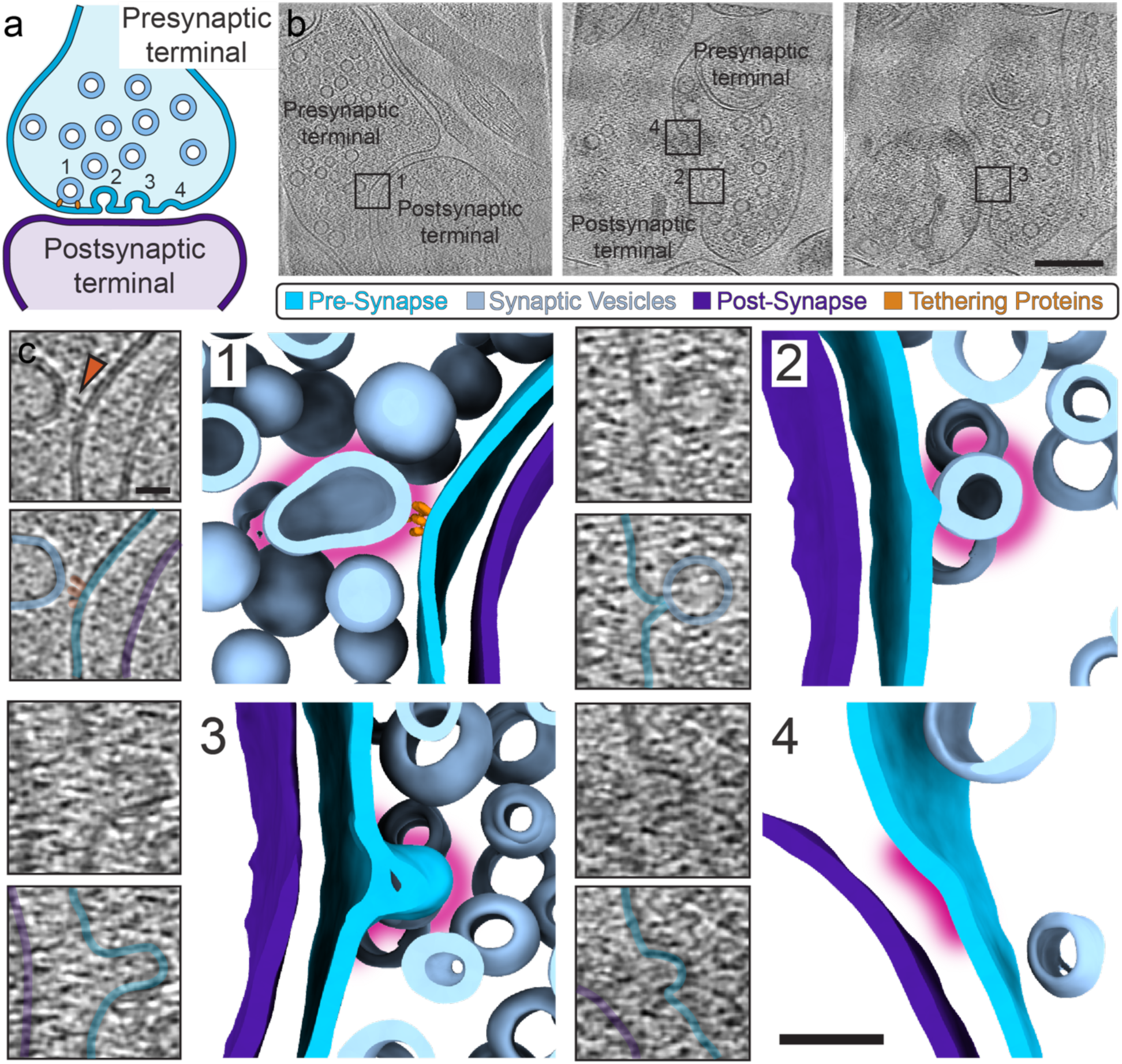
Synaptic Vesicles Fusing with the Presynaptic Membrane. a) Cartoon of synapse highlighting presynaptic terminal and membrane (electric blue), synaptic vesicles (light blue), postsynaptic terminal and membrane (purple) and proteins tethering synaptic vesicles ready for fusion with the presynaptic membrane (orange). b) Slices through two tomograms with synaptic vesicle fusion events. Scalebar 200 nm. c) Zoom in vesicles at different stages of fusion with the presynaptic membrane and corresponding segmentations. Pink halo highlighting the vesicle or curved portion of membrane of interest. Scalebar in tomogram 20 nm, 50 nm in segmentation.

The synaptic cleft could be identified as two closely mated membranes, approximately 20-30 nm^27^ apart with cell adhesion molecules spanning the cleft. The relative populations of molecules on the pre- and postsynaptic membranes could often be different, with molecules on the postsynaptic face not being in contact with molecules on the presynapse, i.e. in some instances, we see these densities continue beyond where the membranes are in contact (Figure 2). In the postsynaptic compartment, we were often able to observe actin filaments. Likewise, we found small clusters of ribosomes only on the postsynaptic side (Extended Data Figure 7), which is consistent with ribosomes being more common in dendrites but not axons. In some tomograms, we could identify a region near the membrane that appears more electron dense, indicative of a putative post synaptic density (PSD). However, importantly, these could not always be observed (Extended Data Figure 7).

### Assessment of the molecular organisation of tissue from the CA1 strata pyramidale to radiatum

One benefit of serial lift-out is that semi-continuous sampling offers insights into structural changes of proximal regions across an organism or sample. By utilising the faster milling rate of the PFIB, a greater amount of material can be removed in a shorter time frame allowing for unique geometries of complex samples to be achieved. When observed from different perspectives, previously unseen aspects of the neuronal organisation can be revealed. To complement our existing data, we developed a strategy to investigate morphological features of the CA1-sr that run within the plane of the sample (Figure 4; Extended Data Figure 10).

**Figure 4.**
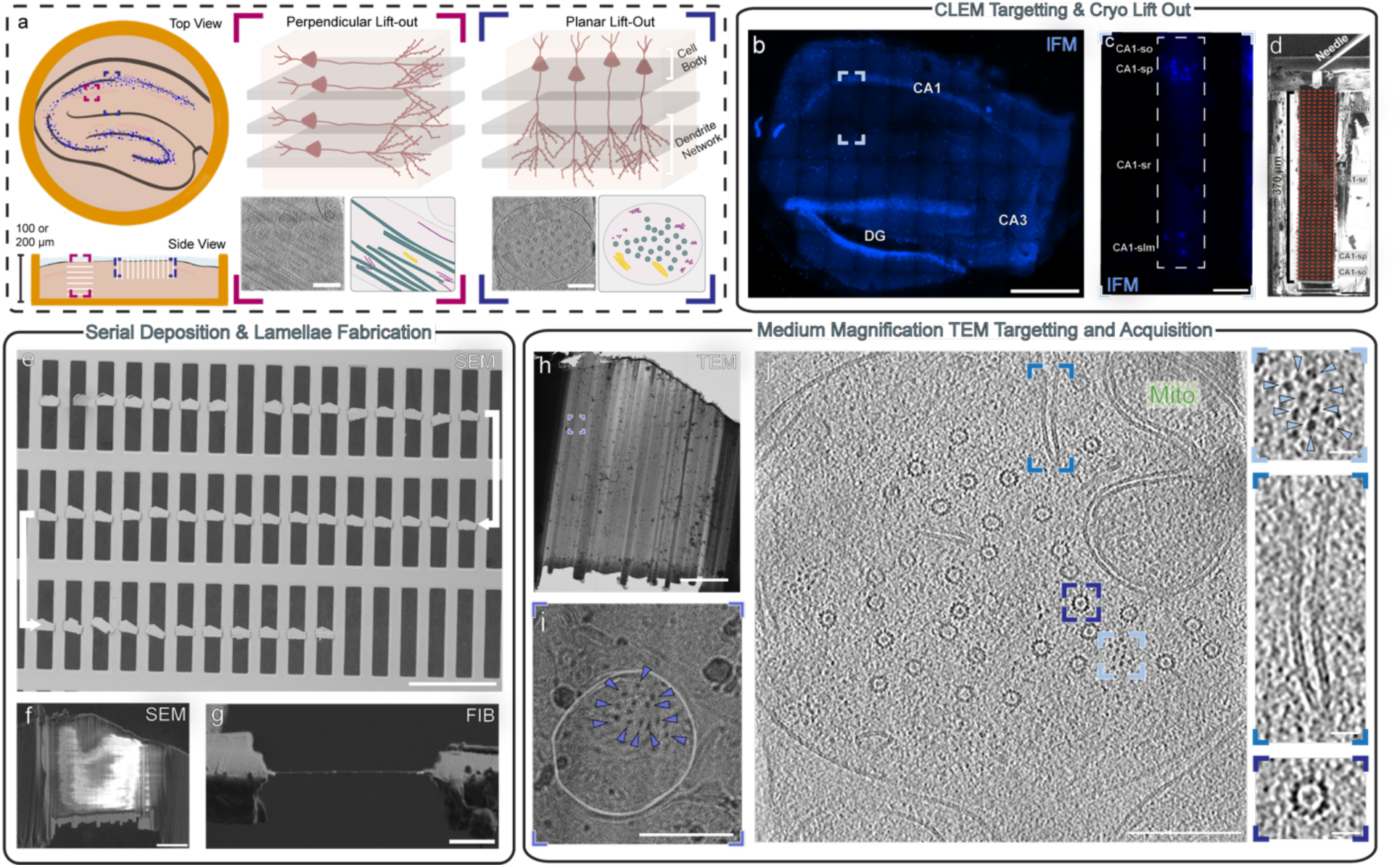
Planar Lift-out of CA1 strata oriens to radiatum. a) Schematic of high pressure frozen hippocampal section with the target region and expected molecular features highlighted. The orientation of the pyramidal cells in Perpendicular (Left) and Planar (Right) lift-out sections are shown, with grey slabs exemplifying serial sections capturing different slices through connected tissue. Examples of tomograms with cellular features annotated highlight the difference appearance in appearance of microtubules (green), endoplasmic reticulum (yellow) and actin (purple) in both lift-out approaches. b) Fluorescence image of a hippocampal section used for planar lift-out with CA1, CA3, and DG marked Scalebar 200 µm. c) Fluorescence image of the lift-out target after trench milling with sub regions labelled. d) FIB image of the 370 µm lift-out target with needle attached and marked with rough locations of serial sections to be sliced off in the next step. Scale bar in d and e 50 nm. e) SEM image of all 42 deposited serial sections. Arrow denotes direction of deposition, starting from CA1-so in the top left through CA1-slm in the bottom right. Scale bar 200 µm. Representative lamella after thinning in the SEM (f), FIB (g), and TEM (h) views. Scalebar f-h is 5 µm. i) Magnified region of (h) shows a dendritic cross section (plasma membrane highlighted in white) with abundant oriented microtubules (blue arrows). Scale bar 1µm. i) Slice through a tomogram with head-on views of microtubules, bundles of actin filaments (blue arrows), extruded endoplasmic reticulum and a mitochondrion highlighted. Scale bar for tomogram 200 nm, scalebars of insets 25 nm.

Using cryo-lift-out, we excised regions that span the transition zone from CA1-sp to CA1-sr to investigate how constituents identified in our samples isolated using the perpendicular approach change with respect to position from sp to sr. To sample across layers of hippocampus, we performed a planar lift out that spanned CA1-stratum oriens (CA1-so) to lacunosum moleculare (CA1-slm) (Figure 4). By mapping the layers of hippocampus using cryo-CLEM as outlined above, a region containing multiple layers of hippocampus was identified and trenches milled using 200 nA Xe around the region of interest (Figure 4b-c). The regions lifted out were 350 and 390 µm in length yielding 40 and 42 sections amenable to thinning of 5 µm thickness, respectively (Figure 4d-e). Serial sections from one of these planar lift-outs were taken forward for lamellae fabrication, where 39 lamellae were thinned from a single, continuous lift out and transferred to the TEM for tilt series acquisition (Figure 4g-h). From 13 imaged lamellae, 246 tomograms were generated spanning a 150 µm region directly below the CA1-sp.

From these data, the cellular composition across the CA1-sr was characterised. Within cells, typical cellular features were identified at each distance across the serial lift out. Features observed include both in plane and perpendicular microtubules (221/246 or 90%), and thinner filaments including actin (75/246 or 30%) and smooth endoplasmic reticulum (ER) (38/246 or 15%) (Figure 4j).

By manipulating the milling geometry and isolating a tissue sample from within the plane of the sample, we could observe CA1 apical dendrites from an axial perspective (25/246) (Figure 4j). Dendrites can be readily identified from TEM search images due to the round plasma membrane and dense arrangement of microtubules within (Figure 4h, i). From this angle, organisation of the dendritic cytoskeleton containing axial microtubules, actin/filamentous networks, and smooth ER could be investigated (Figure 4j, 5). In contrast, perpendicular lift outs in the same region of CA1-sr did not yield many apical dendrite cross sections where head on views of microtubules and actin could be found (1/359), but instead favoured an orientation with side-views of dendrites where groups of microtubules could be found traversing the field of view (10/359) (Extended Data Figure 6b).

### Analysis of the apical dendrite cytoskeleton network from planar lift out

A representative segmentation depicting the axial view of a dendrite illustrates the main cellular components observed across 26 tomograms (Figure 5). The most ubiquitous features were axial microtubules, extruded endoplasmic reticulum, and bundles of actin filaments, found in every tomogram (Figure 5b-d respectively, Extended Data Figure 10a). The number of microtubules observed across the dataset was interrogated in relation to their distance from the CA1-so, ranging from 4-34 per tomogram (Extended Figure 10b). Globally, interfilament distances exhibited a gaussian distribution where nearest neighbours had a mean value of 57.7nm and a median of 54.0 nm (n=481) and exhibited no significant difference as a function of distance across the region (Extended Figure 10c-d). This suggests no trends are evident in microtubule organisation across the regions analysed.

**Figure 5.**
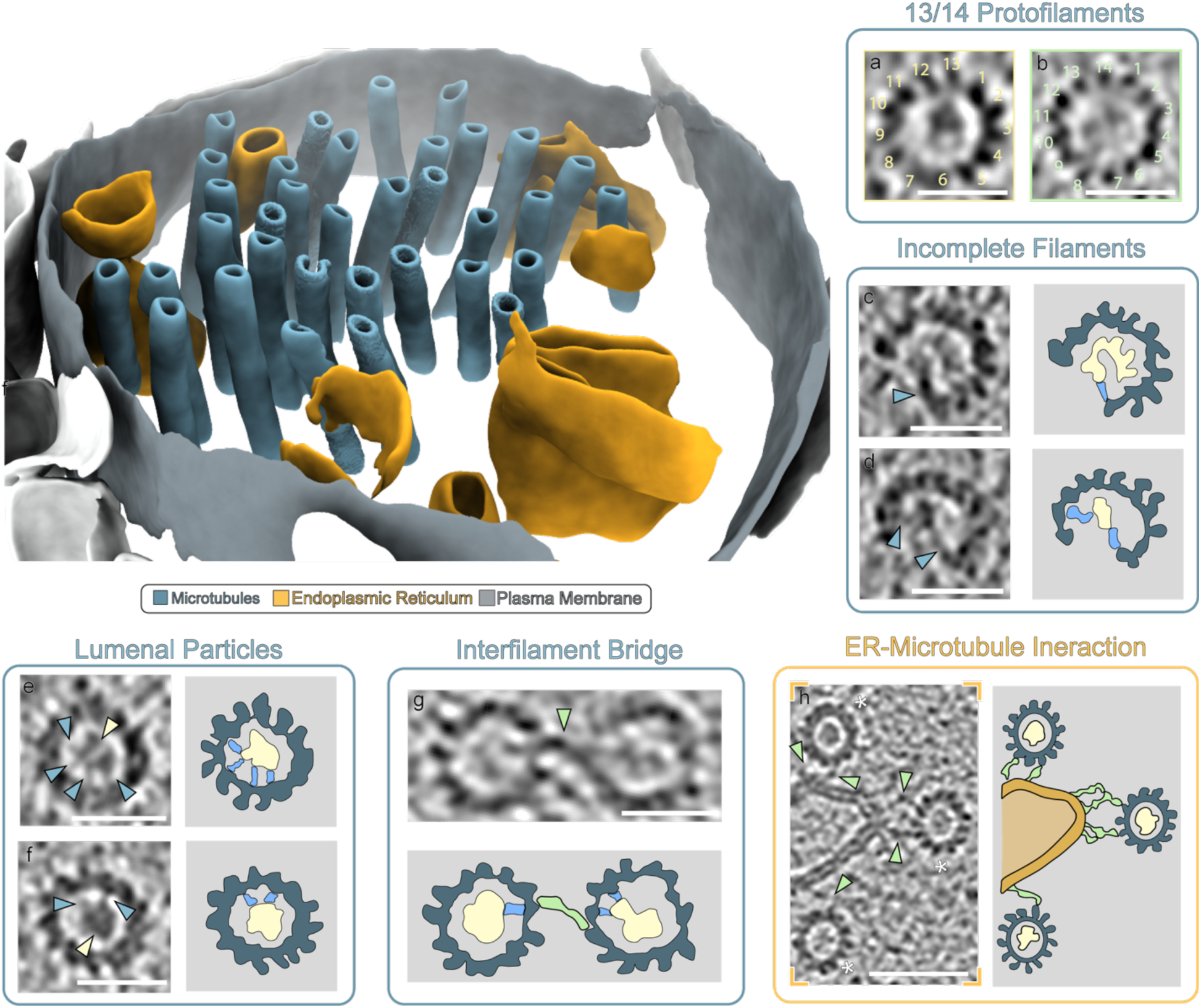
– Characterisationx of the apical dendrite network cytoskeleton from planar lift out. Left) Segmentation of a representative tomogram depicting a cross section of a dendrite with microtubules (blue), endoplasmic reticulum (gold) surrounded by the plasma membrane (grey) highlighted. a-b) Examples of microtubules composed of both 13 pf (yellow box) and 14 pf (green box) respectively. c-d) Examples of an incomplete microtubule, with density seen connecting the edge of the microtubule to an internal density (blue). e-f) Examples of lumenal particles within microtubules (yellow) with multiple associated protein tethers to the inner microtubule wall (blue). g) Example of two microtubules interlinked with a protein tether (green). f) Example of the interactions between microtubules (*) and the endoplasmic reticulum (orange) via protein linkers (green).

Microtubules of both 13 pf and 14 pf could be observed (Figure 5a-b) and had incomplete ends in 3% (15/481) of filaments (Figure 5c-d, Extended Figure 8a). Density was also visible within the lumenal space, with additional density connecting lumenal particles and the inner microtubule wall (Figure 5e-f). Interestingly, bridging densities can be seen between microtubules in proximity, suggesting that at least some microtubules are bundled/cross linked by putative protein tethers (Figure 5g). The microtubules also interact with the ER mediated by multiple protein tether interactions (Figure 5h). Additional, less frequent observations included areas near the plasma membrane devoid of microtubules, displaying a dense, matrix-like structure (9/26 tomograms) (Extended Figure 10a). Mitochondria could also be observed within dendrites (7/26 tomograms), while other double-membraned vesicles of various sizes were seen (8/26 tomograms) (Extended Figure 10a).

## Discussion

Here we demonstrate an approach for the targeting of specific regions of brain tissue for molecular imaging. Notably, this did not require genetically modified animals with fluorescently encoded tags or injection of material, illustrating that with judicious generic labelling, a diverse range of experimental setups and animal models of varying genetic background can be assessed. This can be applied equally with tissues other than brain. Presently, our data represent a benchmark for assessing brain molecular ultrastructure in the CA1-sr within the C57BL6 in-bred mouse.

The use of xenon plasma was particularly advantageous, where large trenches on the order of 500,000 um^3^ of material – 100x the volume of a Hela cell - need to be cleared to isolate material from an HPF carrier using undercuts. Traditional LMIS Ga^+^ FIBs are significantly slower^16,18^, meaning removal of large amounts of material with gallium is impractical. The speed demonstrated here also negates the need for extra cryo-microtomy steps to remove excess material from the tissue and carrier, avoiding additional steps within the workflow^13,31–34^. Consistent with thinning of cellular lamellae^16,17,35^, we were also able to generate thin lamella from tissues using a PFIB (Extended Data Figure 2), demonstrating the versatility of plasma for both clearing large volumes at speed with high currents without sacrificing sample quality during thinning at low currents.

We demonstrate uncompressed preservation of tissue morphology upon vitrification utilising mouse hippocampus as a test case, where hippocampal layers could be easily identified by CLEM. Vitrification of tissues is non trivial, and no thorough assessment of high pressure freezing of tissue exists that demonstrates high quality, vitreous, sample preparation. Therefore, we started from combinations of cryoprotectants that had previously resulted in vitrification of thick specimens^36,37^ while also maintaining tissue health^38^. We incorporated three buffers – phosphate buffer, artificial CSF and an NMDG-substituted artificial CSF^39^ – into our cryoprotectant formulation demonstrating that vitrification can be achieved without removal of maintenance media (Extended Data Table 1). Illustrating the benefit of proper vitrification, our tomograms from vitrified brain tissue contained more open, extracellular space compared to fixed tissue. Similar observations have also been made for tissue that has undergone cryo fixation compared to chemical fixation^22^. Ultimately, this avoids non-native changes in morphology, allowing an investigation of more subtle ultrastructural changes that could underpin developmental or disease mechanisms.

Importantly, in the present work hippocampal slices were maintained at 4°C as samples were prepared. While the time taken for slices to be HPF was 1-4 hours, slices maintained in such a way are viable in organotypic experiments^40^. We did not detect any variation in synaptic vesicle size as a function of post-mortem interval. Nevertheless, the process of HPF and vibratome sectioning will need to be improved in the future for near-pristine preservation from dissection to vitrification be achieved.

Previous work has used high-pressure freezing to generate brain samples for cryo-lift out or FIB milling but are limited to either i) very few (< N=2) examples^14,32,41^ or ii) approaches where only very thin specimens can be frozen^20^. In the former case, there was no assessment of the vitrification while in the latter case, where tissue is frozen “on-grid”, the large forces the brain is exposed to can lead to disruption of tissues during the freezing process, limiting application. In the present study, the method of freezing allows viable tissue slices to be frozen in a format where samples are protected from compression during freezing.

Crucially, the strength is the ability to image molecules within the hippocampus. In our data, we were able to visualise multiple regions and sub-cellular structures. As a focus, we aimed to image in the CA1-sr; in this region of the hippocampus, there is a high degree of pyramidal cells and mossy fibers forming synaptic circuits with neurons emanating from the CA1-sp. Our data are consistent with this, as our data mostly exclude myelin but include multiple examples of active synaptic boutons. The organisation of synaptic contacts within our data show many synaptic vesicles, including membrane fusing with the presynaptic membrane (Figure 3). In some instances, individual synaptic cleft proteins could be visualised (Figure 1, 2). As synapses are larger than the mean free path of an electron (∼ 300 nm), it is not possible to obtain whole synapses within a tomogram, however, the ability to visualise molecules in the synaptic cleft gives indications of the molecular diversity in the synapse.

For our measurements of synaptic vesicles, we used four metrics: diameter, calculated sphere volume from the measured diameter, volume calculated from a convex hull of the segmentation, and sphericity based on the points in the convex hull. For diameter, a large range of vesicle sizes were found, ranging from 41.2 - 51.2 nm as an average per tomogram. We did not find any relationship between vesicle diameter and the sex of the mouse, cryo protectant and buffer composition, post-mortem interval before vitrification, or the day tilt series were collected (Extended Data Figure 8g, Extended Data Table 3). For volume calculated from a convex hull, the missing wedge will obscure the tops and bottoms of vesicles, causing vesicles to have smaller volumes and have lower measures of sphericity.

The ability to extract tissue samples in a variety of orientations allows for lamellae to be fabricated with views previously unobtainable through conventional cellular cryoET experiments. By imaging features from multiple angles, detrimental artefacts arising from the “missing wedge” in Fourier space can be reduced. Furthermore, as the hippocampus is a differentiated tissue, cell populations exist in an organised fashion, where features of interest exist in specific planes. In the present case, dendrites originating from CA1-sp run parallel to one another, traversing our sample plane in the CA1-sr. By lifting out a section of tissue in the plane of the sample, previously inaccessible views of the CA1-sp apical dendrite cytoskeletal network could be imaged in vitrified samples. By preserving the native, hydrated state of the dendrite, fine molecular details that are obfuscated by chemical fixation could be observed (Figure 5). As we integrate multi-scale correlation, the spatial relationship of sub-layers within the tissue with respect to the bulk sample can be maintained and tracked between sequentially deposited sections. Tissue composition, such as cell and synapse types, have been shown to vary from regions of CA1-sr more proximal to CA1-sp to more distal regions^42^. The ability to sample every 5-10 µm facilitates investigation of the subtle changes in molecular composition within and between layers.

In summary, our paper demonstrates a robust, flexible strategy for the preparation of frozen-hydrated, natively preserved (brain) tissue. Importantly, scaling structural biology to tissues requires techniques that can generate enough samples, in a targeted way, that allows a pool of 3-dimensional datasets from cohorts of individuals. This requires reproducibility, high success and feasible timeframes. We were able to generate 10s of tissue chunks per day from individual slices, which incorporate large-scale mapping from micron to nanometre scale. Our data demonstrate pristinely preserved molecular information, allowing an analysis of molecular context within brain structures. This lays a framework for the routine assessment of tissue samples in the future, where specific features relating to pathology may be correlated and assessed on a molecular scale.

Crucially, this allows an assessment from different samples on the order of days, potentially enabling targeted clinical observations at a significant scale. Future work will incorporate the integration of scanning electron methods as well as machine-learning to improve automation and feature recognition.

## Methods

### Animal Handling and Brain Dissection

All animal experiments were monitored and facilitated by Named Animal Care and Welfare Officers from the Mary Lyon Centre for Mouse Genetics at MRC Harwell, and animals were treated in accordance with the UK Animal Scientific Procedures Act (1986). C57BL/6J mice (Jackson Labs, Bar Harbor, ME) were provided by the Mary Lyon Centre for all datasets collected.

Adult mice were euthanised in a schedule 1 procedure via overdose with an intraperitoneal injection of dilute pentobarbital in buffered saline solution (1:1). Death was confirmed through permanent cessation of circulation via cutting of the femoral artery. In total 11 mice were used in this study. Acute hippocampal slices for cryo-ET evaluation of the CA1 architecture were prepared from one male and one female 4–6-month-old C57BL/6J mouse. For the cortex experiment datasets, a 5-month-old female C57BL/6J mouse was used. When assessing vitrification and optimising early lift-out methodologies, samples from mice aged 7 to 184-days-old (male and female) were high pressure frozen. Mouse pups (between ages of 7-21-days-old) were euthanised as stated previously but instead using an intraperitoneal injection of 200 mgmL^-1^ concentrated pentobarbital.

### Sectioning of Acute Brain Slices

The slice preparation protocol was modified from established protocols^43–45^. In brief, following brain dissection from the mice, the cerebellum was removed and the hemispheres separated at the longitudinal fissure using a scalpel. The remaining tissue was glued cut face down onto a chuck, and 100, 150, or 200 μm sagittal slices were made using a Leica VT 1200S Vibratome set at a slicing speed of 0.7 mm/s and an amplitude of 1 mm in ice-cold dissection medium (99% HBSS, 0.035% ascorbic acid, 0.002% ATP, 1% penicillin/streptomycin solution, filter-sterilised through a 0.22-µm membrane, pH 7.1). Two brains were sectioned while bubbling with carbogen (95% O_2_/5% CO_2_): the cortex where data are presented and datasets from the 172 day old female mouse (“CA1-sr 2” and “CA1-sr 4”). Where cortex is used, biopsy punches were taken directly from sagittal sections. For hippocampus biopsies, each hippocampus was first isolated using curved tip teasing needles (Bochem) and individual hippocampi transferred to a well of a 24-well plate with resting media (300 µL Hibernate-A media (Gibco) supplemented with 2% B-27 (Gibco)) for recovery and transport until high pressure frozen.

### High Pressure Freezing

3 mm type B gold-plated copper high pressure freezing carriers (Leica Microsystems) were prepared as previously described^19^. Briefly, the flat side was sanded with 4000 grit sandpaper to remove machining marks followed by 10000 grit sandpaper to remove harsh aberrations followed by metal polish to smooth the surface. Sanded and polished type B carriers and type A carriers were incubated in hexadecene (Sigma-Aldrich) for at least 45 minutes prior to use. Type A carriers containing tissue were pen-marked on the rim with to aid with carrier orientation for subsequent imaging and serial lift-out.

Tissue slices in resting media were incubated with Hoechst 33342 (Invitrogen) nuclear stain for 5 minutes prior to target excision with a 2 mm biopsy punch. Biopsies were transferred via transfer pipet to a 24 well plate with cryoprotectant in buffer (Extended Data Table 2) where they were allowed to incubate at room temperature for the specified time ranging from 0-30 minutes. After incubation, tissue biopsies were transferred to the 0.1 mm (100 µm thick tissue sections) or 0.2 mm (150 or 200 µm thick tissue sections) recessed side of 3 mm Type A gold-plated copper HPF carriers.

In total, brain dissection, vibratome slicing, and hippocampal isolation from all slices took approximately 1 hour, followed by approximately 20 minutes of slice incubation with cryo protectant to allow for complete vitrification. Each slice took several minutes to manipulate and HPF, and only slices which appeared optimal in both fluorescence and the FIB/SEM were used for subsequent lift-out and data collection. This resulted in postmortem intervals for slices used for data collection ranging from 1.5 - 4 hours.

### Cryo-confocal Microscopy

After freezing, carriers were screened on a Stellaris 8 Cryo Confocal Microscope (Leica Microsystems) fitted with a cryostage. Imaging in fluorescence and reflection modes were performed in camera mode with an HC PL APO 50x 0.9-NA objective using the LAS X software (Leica Microsystems). Tilesets were acquired with 20% overlap between tiles. Tilesets were merged and maximum intensity projections calculated in the LAS X software for fluorescence and reflection modes. An outline of the tissue section was often visible in reflection mode and used in combination with the carrier rim to aid in alignment between reflection and fluorescence channels. The mark drawn onto the carrier rim was visible in reflection mode and used for orientation-conscious carrier loading into the FIB/SEM.

### Hippocampal Layer Targeting

Carriers were oriented for loading into the Helios “G5” Hydra CX plasma FIB/SEM (Thermo Fisher Scientific) using previously obtained information from the cryo-confocal microscope about mark location relative to hippocampal layer orientation. When orientation information was available, samples were loaded into the shuttle such that the CA1-sp would face down with the dentate gyrus facing up. This would allow the CA1-sp to be oriented up in the fluorescence module and the long trench required for perpendicular lift-out to stretch into the CA1-slm such that lift-outs could be obtained from both the CA1-sr and the directly overlaying CA1-sp. This orientation maximized the number of lift-outs that could be obtained from these layers of interest.

Approximately five uniquely shaped fiducial markers were milled into the tissue sample surface that would be visible in the SEM and the integrated fluorescence module (IFM). These were composed of 75 µm x 4 µm x 3 µm Z depth (Si) rectangle patterns milled at 60 nA with xenon plasma. One fiducial was placed near the center of the carrier to aid in centering the carrier in the IFM for fluorescence tileset acquisition.

A Meteor fluorescence module (Delmic) incorporated onto the Helios Hydra FIB/SEM and equipped with a 20x (NA 0.45) objective was used to acquire tilesets that would span the entire carrier in X and Y. This was 11 x 9 tiles (1.88 mm x 1.87 mm) with 10 Z steps of 2 µm per step. Laser power was set to 500 mW and 150 ms exposures were acquired. Maximum intensity projections were calculated in the Odemis software (Delmic).

Fiducials visible in the fluorescence image were used as reference points for targeting specific hippocampal layers. If fiducials were far from the target region, smaller (30 µm long line patterns with 3 µm Z depth) were milled using 4 nA xenon plasma current adjacent to the lift-out target region and checked in the fluorescence module before trench milling to ensure fine targeting. For CA1-sr, targets were placed up to halfway between the CA1-sp and the visible CA1-slm (Figure 1). At closest to the pyramidale layer nuclei this was 50 um and 150 um at farthest.

### Serial Lift-Out

Serial lift-out was performed on a Helios “G5” Hydra CX plasma FIB/SEM (Thermo Fisher Scientific) equipped with a tungsten EasyLift needle. A copper block of 15 (width) by 10 (depth) by 12 (height) µm was taken from the receiver grid and attached to the end of the needle by redeposition welding^21^.

For perpendicular lift-out, a target area of 60 µm in X and 30 µm in Y was chosen based on position relative to layers observed in the fluorescence module. The stage was compucentrically rotated such that the sample would be perpendicular to the FIB. For our stage with a 27° pre-tilt sample shuttle, this was a stage tilt of 25°. A long trench of 60 µm in X by 150-200 µm in Y by 5-6 µm in Z was milled with an RCS pattern behind the target region such that the long trench would be at the front of the sample after compucentric rotation with scan rotation set to 180° (Extended Data Figure 1a, Extended Data Table 2). A second, shorter, but deeper trench (RCS pattern) was milled in front of the sample to allow for subsequent assessment of completeness of side and undercuts in the next step. This milling pattern was 60 µm in X by 40-60 µm in Y by 6-11 µm in Z. Larger Z depths were used in instances where longer trenches were also used (Extended Data Table 2). The scan direction of both trench RCS patterns were oriented towards the lift-out target area. Side and undercuts were made with the long trench facing towards the FIB at a stage rotation of 8-15 degrees with our 27° shuttle. For side cuts (15 nA), patterns were 4 µm wide with at least 45 µm between patterns. For undercuts, patterns were 6-8 µm tall (Extended Data Figure 1b).

For sample attachment to the needle, the stage was rotated to approximately 0° - or the shallowest angle where the entirety of the remaining tabs leftover from the side cuts could be seen. The tissue was then attached to the copper block adapter on the EasyLift needle by redeposition welding (Extended Data Figure 1c). Once attached, the remaining tabs on the tissue were milled away with 4 µm wide rectangle milling patterns at 4 nA. The EasyLift needle with attached tissue sample was then retracted.

For planar lift-out a target area of 60 µm in X and 350–400 µm in Y was chosen based on position relative to layers observed in the fluorescence module. The stage was compucentrically rotated such that the sample would be perpendicular to the FIB. For our stage with a 27° pre-tilt sample shuttle, this was a stage tilt of 25°. Two trenches of 50 µm in X, 350–400 µm in Y and 6 µm in Z were milled either side of the region of interest using 200 nA. A third trench was milled at the base of the region measuring 210 µm in X, 60 µm in Y and 2 µm in Z which is where the lift-out needle with the copper block adapter will be attached (Extended Data Table 2). This left the region connected to the bulk of the material The scan direction of both trench RCS patterns was oriented towards the lift-out target area. Undercuts were performed at ±90° relative to the region of interest, with a −15° stage tilt. Rectangular patterns that span the width of the lift out region (350–400 µm) in X, 10 µm in Y and 1 µm in Z were placed 10 µm below the surface of the sample. Milling was performed at 15 nA until no material connected the region and the bulk of the sample, confirmed by SEM/FIB imaging.

For sample attachment to the needle, the stage was rotated perpendicular to the FIB. For our stage with a 27° pre-tilt sample shuttle, this was a stage tilt of 25°. The tissue was then attached to the copper block adapter on the EasyLift needle by redeposition welding (Extended Data Figure 1c). Once attached, the material connecting the side of the region and the bulk sample were milled away with 5 µm wide rectangle milling patterns at 4 nA. The EasyLift needle with attached tissue sample was then retracted.

### Serial Section Deposition

Serial sections were deposited onto rectangular pattern 400 x 100 mesh TEM support grids (Agar Scientific). The uneven bottom portion of the lift-out chunk was milled away before section deposition and any extraneous material at the sides wider than the top lip on the grid bars was trimmed away at 4 nA. The stage was rotated to the shallowest angle possible, here –5 – −3°, before the lift-out chunk was brought down into contact with the grid. 3-5 µm sections were then milled off using a line pattern at 1-4 nA. While 4 nA line patterns were faster (<2 min / section), 1 nA line patterns (∼5 min / section) yielded a smoother surface that would be advantageous for subsequent thinning steps, thus 1 nA was used for sectioning in most cases. After all sections had been deposited, the stage was rotated to 15° and welding patterns were placed on either side of each section for attachment to the grid by redeposition welding. These patterns were CCS oriented towards the copper bars with a 30 us dwell time. Patterns were 3 µm long by 0.8 µm tall with approximately 4 µm periodicity. This resulted in 6-8 welds per section and took ∼30 s/weld. After all sections were deposited and welded, GIS was applied for 70 seconds to achieve a few hundred nm thick GIS layer.

### Fine Milling of Serial Lift-Out Sections

Thinning of serial sections was carried out on an Arctis plasma FIB (Thermo Fisher Scientific) initially using AutoTEM (Thermo Fisher Scientific) for automated thinning with xenon plasma down to approximately 400-600 nm thickness depending on section quality where higher quality sections could be automatically thinned to lower values. Lamella width was set to 15-18 µm with a target thickness of 110 nm and a Z depth of 2.5 µm in silicon. For rough milling, rectangle patterns with a beam current of 4 nA with 2-3 µm pattern offsets were used to bring the total lamella thickness down to ∼4 - 6 um. For medium milling, a cleaning cross section was used with 0.7-1.0 µm offsets with a beam current of 1 nA to bring the lamella thickness down to ∼1.4–2 µm. For fine milling, cleaning cross section patterns with 120 – 300 nm offsets were placed and a beam current of 0.1 nA was used to mill sections down 400-600 nm before switching to manual polishing steps with argon plasma to bring the final thickness down to ∼150-300 nm using 60 pA and 20 pA beam currents (Figure 2d). For all automated milling steps, we found that the lower end of the thickness spectrum given at each step could be used for high quality, smooth, stable starting sections while values at the higher end of the range were needed for lower quality sections starting with uneven surfaces before moving to manual polishing with argon plasma.

### Cryo-Electron Tomography Tilt Series Acquisition

TEM image acquisition was carried out on a Titan Krios G4 (Thermo Fisher Scientific) electron microscope operating at 300kV and equipped with a ± 90° stage, a Selectris Energy Filter (Thermo Fisher Scientific), and a Falcon 4i direct electron detector camera (Thermo Fisher Scientific). For all acquisitions, an energy selecting slit width of 10 eV was used. Dose symmetric tilt series were collected utilising an image-shift/beam-shift data collection strategy in Tomo5 version 5.17.0.6390 (Thermo Fisher Scientific). Tilt series were acquired as movies using the .EER file format at a nominal magnification of 42,000x (3.05 Å/pixel) or 64,000x (1.98 Å/pixel) in counting mode from +60 to –60° starting from the predetermined milling angle (i.e. zero degree offset) in 3° increments with a total dose of 130 e^-^/Å^2^ with a target defocus from –3 to –5 µm.

### Tomogram Reconstruction

Warp^46^ version 1.0.9 was used for CTF estimation and motion correction of tilt series. Tilt series were aligned and reconstructed using AreTomo^47^ version 1.3.4 and tomograms were visualized and lamella thickness measured using IMOD version 4.12.56. Thickness measurements were based on selecting a position near the center of the tomogram, moving through Z until biological material was no longer visible and setting that as the starting point of the lamella. This was repeated moving through Z in the opposing direction where the distance between the two points was the measured lamella thickness. This measured thickness was then used to reconstruct tomograms again with a more accurate AlignZ parameter. Bin8 tomograms (pixel size 15.84 Å) were post-processed using Isonet^48^. Tomograms were subjected to CTF deconvolution (SNR fall off = 0.9, deconv_strength = 0.9 – 1.1 depending on dataset) before missing wedge correction. All tomograms shown in figures are deconvolved, but not missing wedge corrected.

### Tomogram Segmentation

Membrain-seg^49,50^ was used for initial segmentation of tomograms post-processed in isonet. Besides membranes, membrain-seg also segmented high contrast structures like microtubules. Membranes were further segmented using Chimera’s segger tool. Microtubules and the synaptic cleft were segmented manually in Amira version 2023.1.1 (Thermo Scientific). Amira was also used to manually segment some regions of membrane and synaptic vesicles not segmented with membrain-seg and to segment vesicle tethering proteins. For manual segmentations, a 20 Å gaussian filter was used to aid in segmentation of densities within microtubules and smoothing of membranes.

The package Volume Segmantics^51^ was used for quantitative segmentation of synaptic vesicles. The training dataset was comprised of 22 tomograms and their corresponding binary masks, which were generated through a combination of manual annotations using Napari^52^ and training intermediate models and correcting their predictions (pseudo-labelling). Different model architectures were tested to optimize the model, ultimately using a U-Net with ResNet50 encoder which was trained for a total of 13 epochs. A combination of 0.75 Binary Cross Entropy and 0.25 Dice Loss (BCEDiceLoss) was used. The segmentations were post-processed by applying a threshold for sphericity and minimum voxel size to exclude any partially segmented vesicles. Python scripts were developed to calculate key morphological metrics, such as the sphericity, volume and maximum diameter of each synaptic vesicle. These scripts utilized the NumPy^53^, SciPy^54^ and Scikit-image^55^ libraries, with the volume measured by computing the convex hull of each synaptic vesicle using the ConvexHull function from the scipy.spatial module. This provided the smallest convex shape enclosing all points of each vesicle. The surface area of the convex hull was also obtained, and these values were used to calculate the sphericity of each vesicle. We note that the missing wedge will result in a smaller convex hull volume than the calculated sphere volume from the diameter and a lower calculated sphericity than may exist in reality.

### Filament Analysis

Microtubules backbones were traced through low-pass filtered tomograms to create models in IMOD^56^ (C) according to a previously published protocol^57^. Model files were converted into coordinates where analysis and visualisation were carried out using custom python scripts.

## Contributions

M.G. conceptualized the work. R.C. and M.E.S. performed vibratome slicing and hippocampal dissection. C.G. and J.L.R.S. carried out high pressure freezing. J.L.R.S. and C.G. optimised the initial lift out with input from R.C. and T.S.G. C.G. collected cryo confocal images. M.C. and C.G. developed the targeting approach for lift-out. C.G. and J.L.R.S. carried out serial lift-out and FIB/SEM experiments. C.G., M.C., and J.L.R.S. collected cryoET data. C.G. and J.L.R.S. reconstructed tomograms. A.K. and A.P. performed segmentation and analysis of synaptic vesicles. C.G., J.L.R.S., and M.G. analysed the data. Initial draft was written by C.G. and J.L.R.S. and edited by C.G., J.L.R.S., and M.G. with input from all authors. Funding was acquired by M.G.

## Acknowledgements

We would like to thank Dr. Sara Wells, Dr. Marianne Yon, Dr. Michelle Stewart, and Jessica Podd from the Mary Lyon Centre for Mouse Genetics at MRC Harwell for their support in animal work. We would like to thank Dr. Victoria Garcia-Giner for assistance acquiring cryo-confocal images. We would also like to thank Dr. Casper Berger, Dr. Charlie Lovatt and Helena Watson for their assistance with data processing and analysis. This work was supported by a Wellcome Trust Development Award (225902/Z/22/Z to M.G.) and the Electrifying Life Science project (220526/Z/20/Z to Dr. James H. Naismith). J.L.R.S. is supported by a Wellcome Trust PhD Studentship (226810/Z/22/Z). The Rosalind Franklin Institute is funded by UK Research and Innovation through the Engineering and Physical Sciences Research Council (EPSRC).

## Data Availability

The raw cryoET data used in this paper can be found on the EMPIAR data repository (https://www.ebi.ac.uk/empiar/) at the following accession code: EMPIAR-XXXXX.

