## Extended data figures for "Charting the molecular landscape of neuronal organisation within the hippocampus using cryo electron tomography"

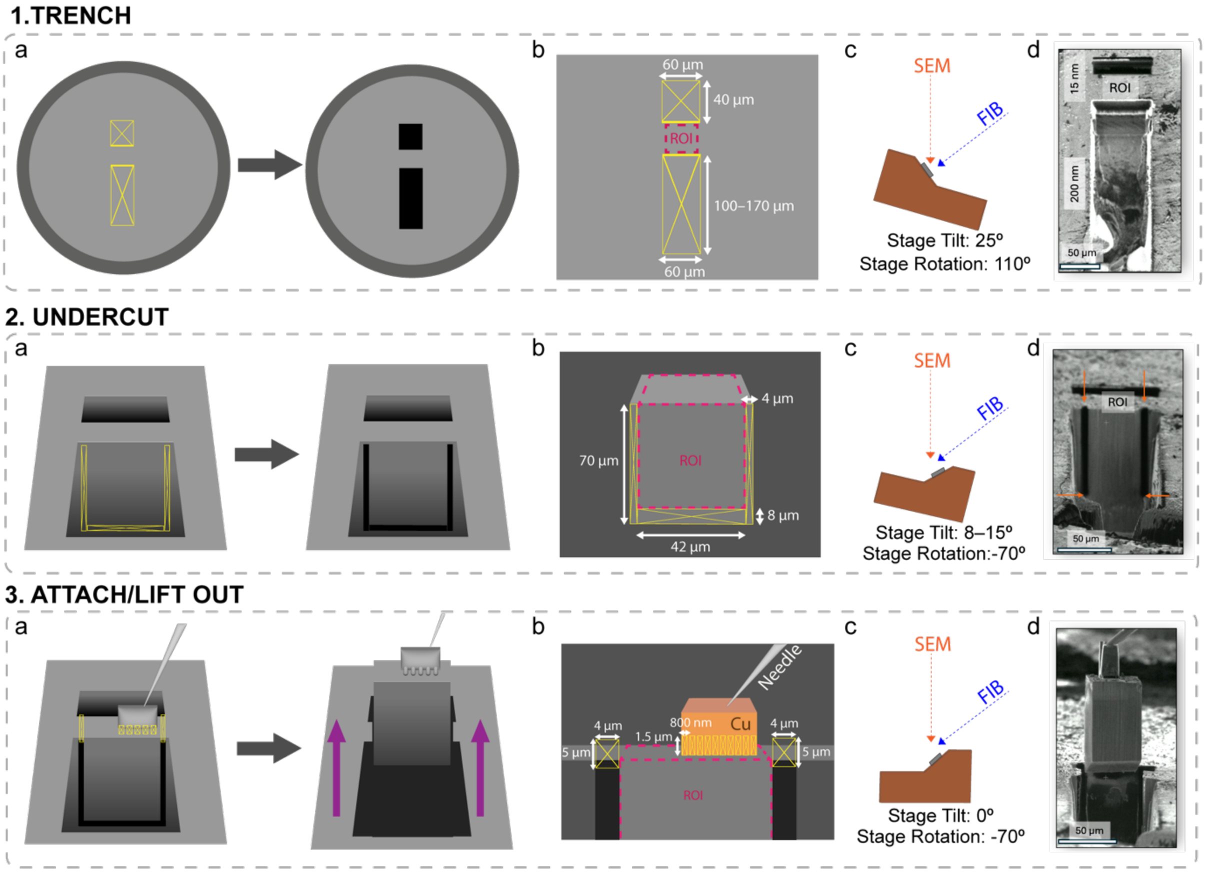


**Extended Data Figure 1. Milling Procedure for Cryo-Lift Out of Perpendicular Samples from HPF Carriers.** 1a, 2a & 3a) Schematic diagram showing the FIB milling patterns (yellow) performed on the sample for Trench Milling, Undercutting and Attachment & Lift Out steps respectively. For the Attachment & Lift Out step, the lift out needle is depicted on top of the region of interest and the purple arrow denotes the direction of the lift out. 1b, 2b & 2c) The region of interest (ROI) (Pink) and the dimensions of FIB milling patterns (yellow) placed for Trench Milling, Undercutting and Attachment & Lift Out steps respectively. 1c, 2c & 3c) Schematic diagram for the orientation of the SEM and FIB beams in relation to the FIB sample holder, with stage tilts and rotation angles for Trench Milling, Undercutting and Attachment & Lift Out steps respectively. 1d, 2d and 3d) Examples of FIB images for Trench Milling, Undercutting and Attachment & Lift Out steps respectively.

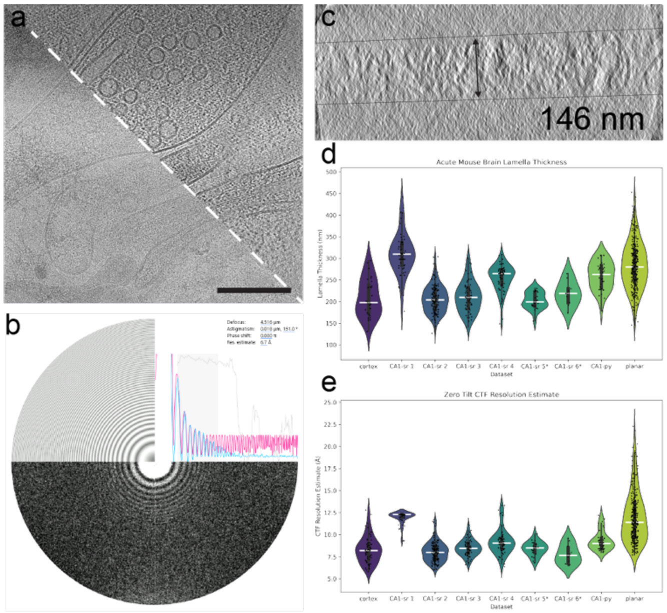


**Extended Data Figure 2. Quality of CryoET data from lifted-out mouse brain**. a) Raw 0º tilt image before preprocessing motion correction (bottom left) and after reconstruction (top right). Scale bar 200 nm. b) Contrast transfer function experimental (blue) and estimated (pink) functions, defocus, and resolution estimates calculated in the Warp software package[46]. c) Reconstructed tomogram thickness measured in IMOD. d) Thickness measurements from all tomograms in the 9 vitreous datasets acquired in this work. The cryo protectant used for all datasets was kept consistent, except for datasets with a *, where CA1-sr 5 was frozen in 10% Dextran, 10% Sucrose in aCSF, pH 7.4 and CA1-sr 6 was frozen in 10% Dextran, 5% Sucrose, 5% Ethylene Glycol in aCSF, pH 7.4. e) CTF Resolution Estimate from the 0º tilt output from Warp during initial image processing.

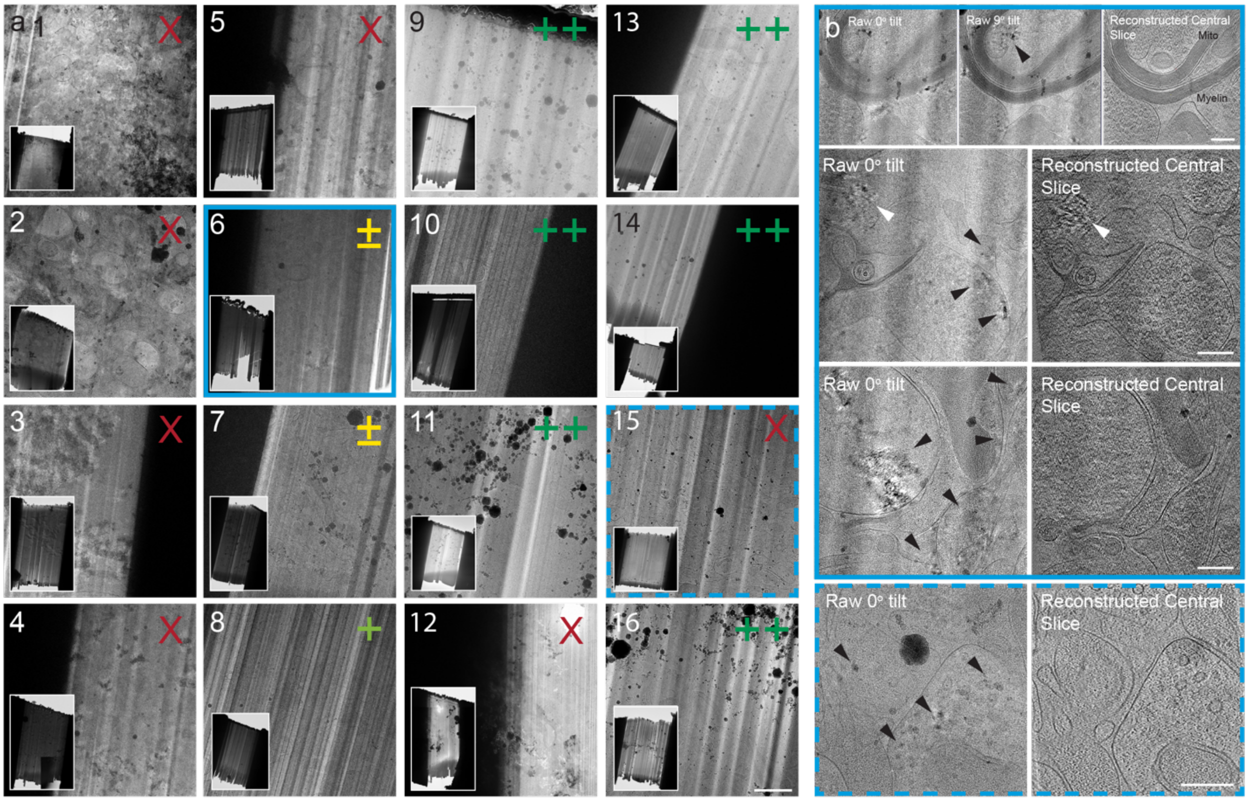


**Extended Data Figure 3. Vitrification Screen for Acute Mouse Brain Tissue Slices**. a) TEM of representative whole lamella overviews (left) and higher magnification search maps (right) from each tissue slice thickness, cryo protectant, buffer, and incubation time combination detailed in Extended Data Table 1. ”X”, ”±”, ”+” and ”++” indicate degree of vitrification. An ”X” represents conditions where no areas of the lamella appeared vitreous and tilt series were not able to be collected without stage tracking areas resulting from extreme fluctuations in contrast introduced by ice diffraction. A ”±” indicates lamella where tilt series could be acquired, but frames in all tilt series displayed incomplete vitrification evidenced by ice diffraction. A ”+” rating was given to conditions where some tilt series could be acquired without ice diffraction, but some non-vitreous ice could be seen in some tilt series. A ”++” symbol was assigned to conditions where no or very few (<10%) tilt series displayed evidence of non vitreous ice. Scale bar 2 µm. b) representative raw tilts from condition 6 (top, solid outline) and condition 15 (bottom, dashed outline) illustrating that while signs of non-vitreous ice may (white arrows) or may not (black arrows) be present in tomogram reconstructions, raw tilts often displayed signed of non vitreous ice. Scale bars 200 nm.

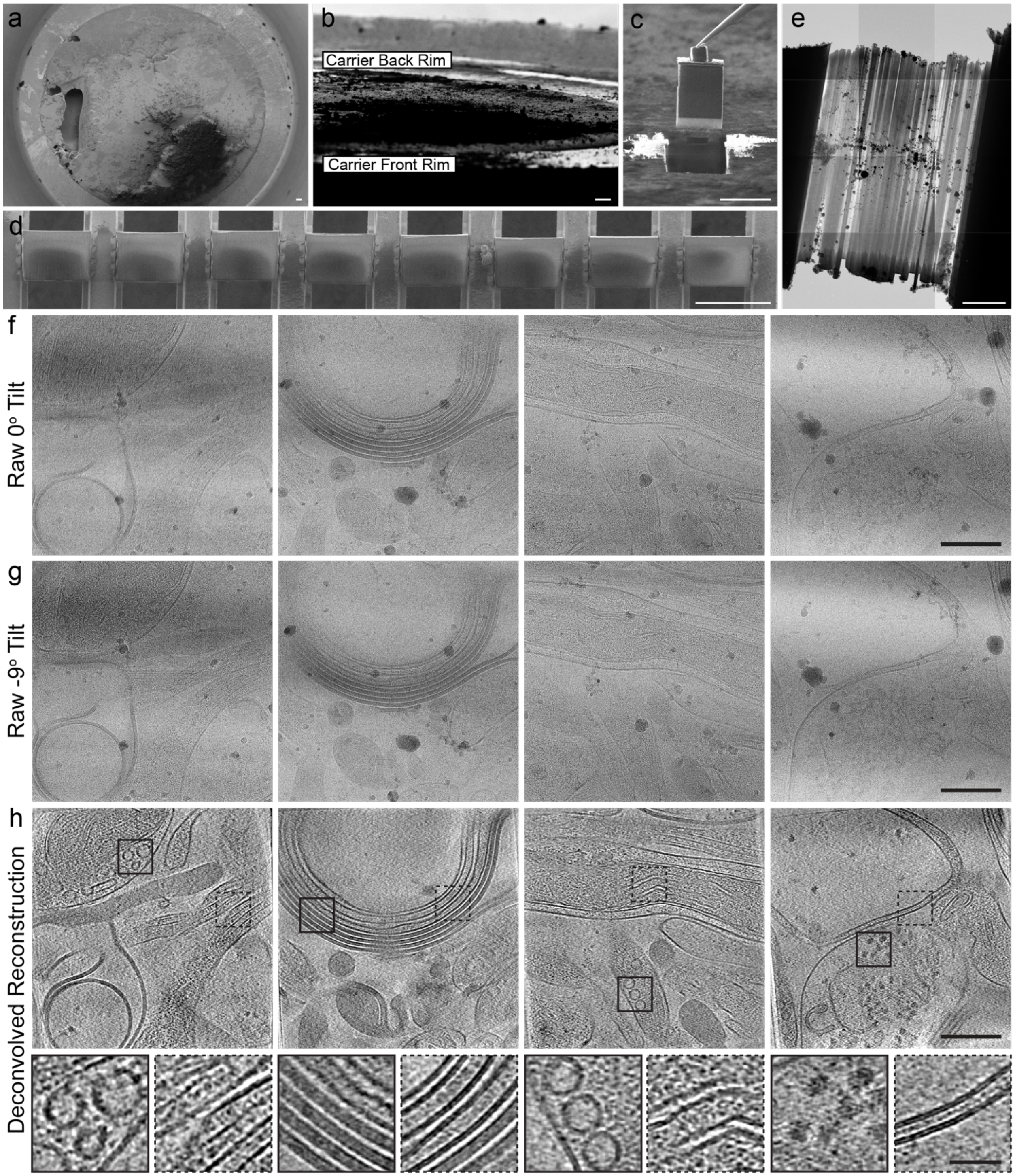


**Extended Data Figure 4. CryoET of Vitrified Mouse Cortex**. a) SEM image of a high pressure frozen biopsy of mouse cortex vitrified in a high pressure freezing (HPF) carrier. b) FIB view of cortex sample in HPF carrier. c) lift-out and d) serial sections deposited to TEM grid. Scale bar a-d 50 µm. e) Thinned lamella (scale bar 5 µm) with raw tilts taken at 0 degrees relative to the milling angle (f) and -9 degrees relative to the milling angle showing no diffraction from non vitreous ice or membrane deformation indicative on a non vitreous sample. h) Deconvolved reconstructions highlighting vesicles, microtubules, myelin, mitochondrial cristae, and ribosomes.  Scale bars f-h 200 nm, insets in (f) 50 nm.

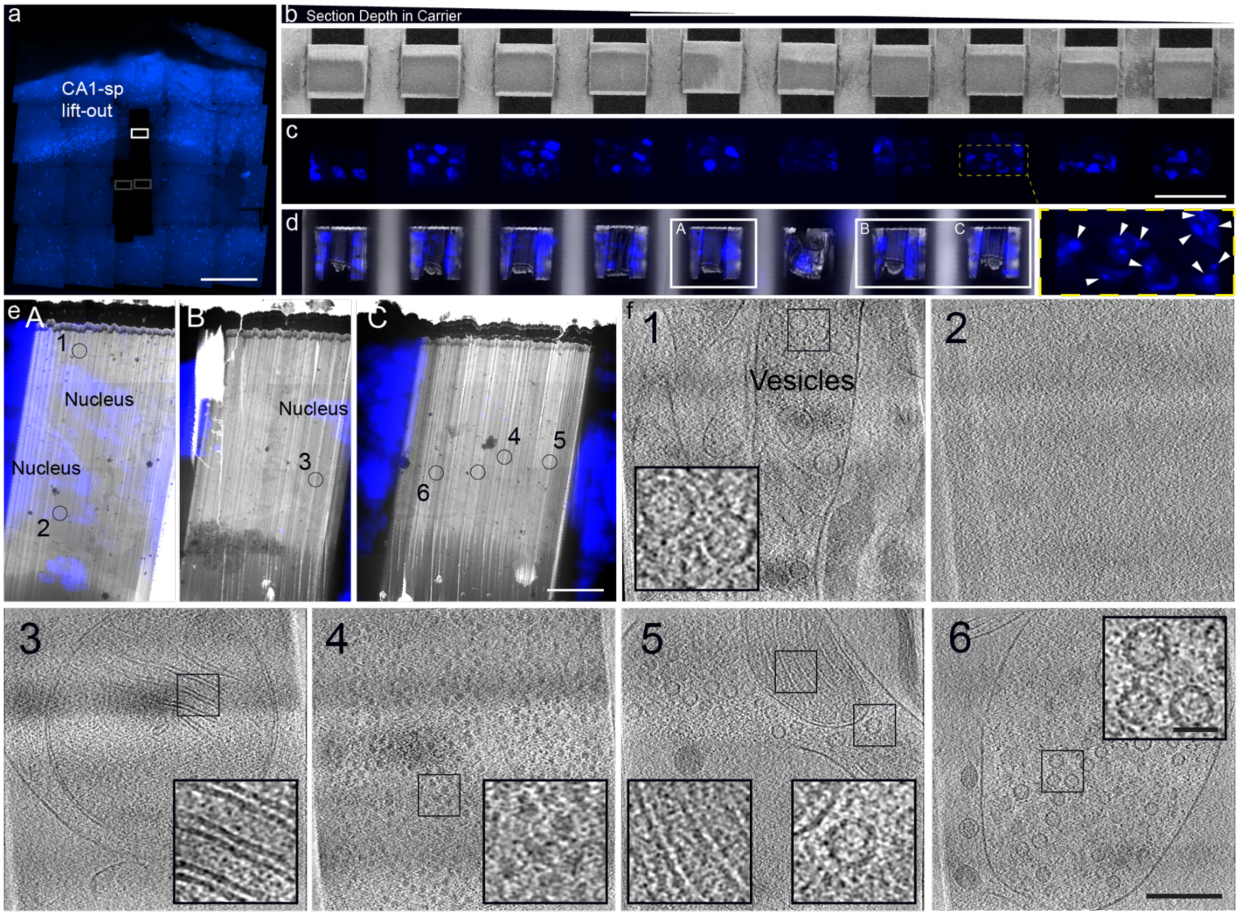


**Extended Data Figure 5. Fluorescence targeting and features in CA1-sp.** a) Fluorescence tileset of the same carrier used in Figure 1 showing the lift-out positions for CA1-sr (gray boxes) and CA1-sp (white box). Scale bar 200 µm. SEM (b) and fluorescence (c) images of serially deposited sections from the CA1-sp lift-out. Scale bar 50 µm. Inset in c) highlighting diffuse fluorescence indicative of euchromatin and more punctate fluorescence (arrows) highlighting heterochromatin. d) Thinned lamella with fluorescence overlay in the SEM with blown up versions of A-C where fluorescence and TEM overviews are overlaid and select tilt series acquisition positions marked (e). Scale bar 5 µm. f) Slices through reconstructed tomograms acquired at positions 1-6. Position 2 was acquired in a fluorescent, nuclear, region of the lamella. Insets highlight vesicles, mitochondria cristae, and ribosomes. Scale bar for tomograms 200 nm, 40 nm for insets.

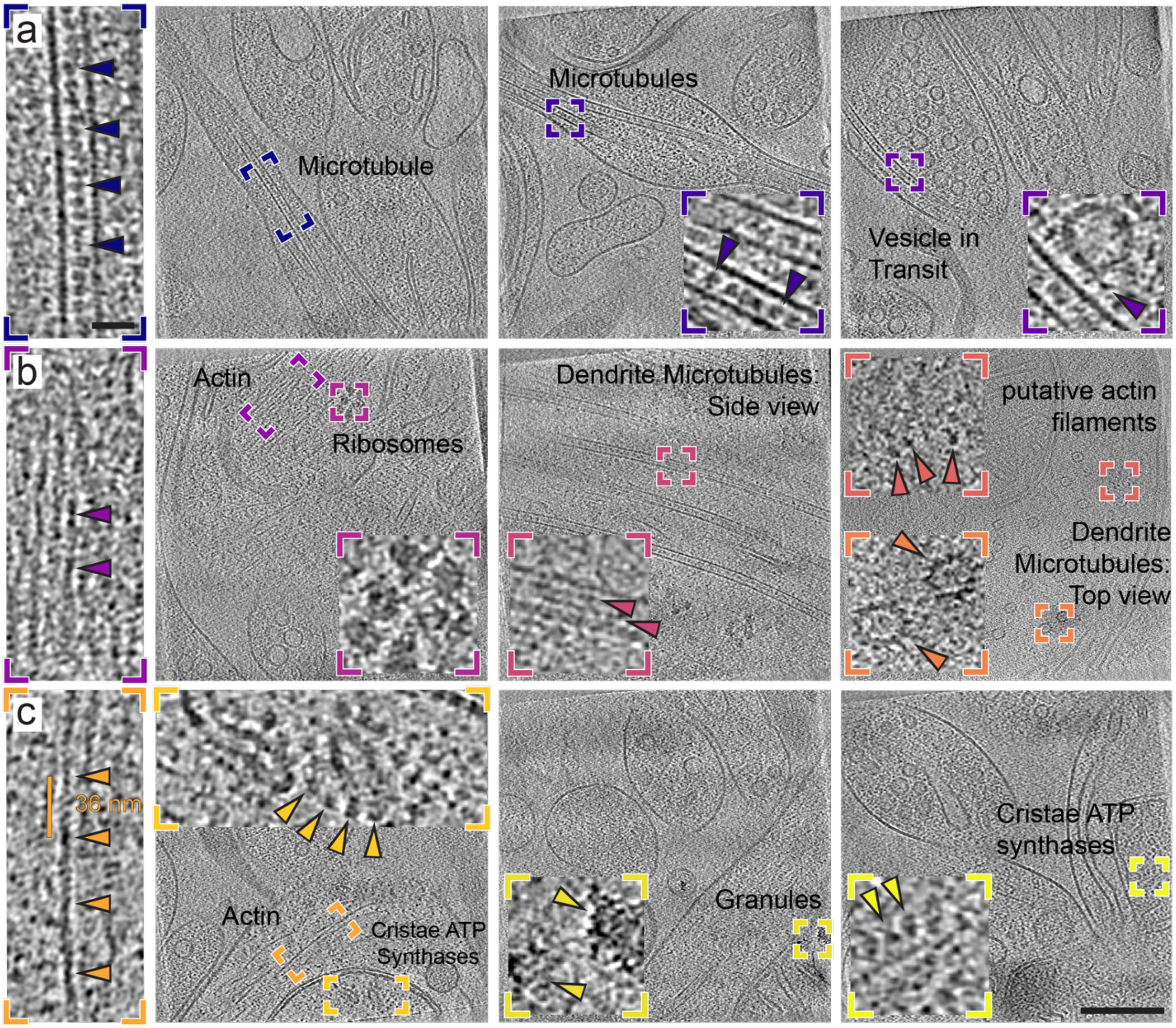


**Extended Data Figure 6. Features Observed in CA1-sr**. In perpendicular lift-outs from the CA1-sr we were able find abundant microtubules with luminal bodies (a, left and middle arrows) and in rare instances vesicles being transported along microtubules (right). We also less commonly found larger cytoskeletal organisation (b) in the form of actin filaments (left), dendrites with microtubules traversing the entire field of view (middle, 10/359 tomograms) and very rarely head on views of dendrites with microtubule (right, bottom inset, 1/359 tomograms) and putative actin filament cross sections (right, top inset). For traversing microtubules, we could occasionally observe the lattice (arrows in middle inset) of aligned subunits. Along with actin, whose identification was supported by a 36 nm helical repeat (c, leftmost inset) we could independently find mitochondria with (middle, 13/359 tomograms) and without (left, right, 119/359 tomograms) granular deposits. ATPases could be observed in cristae (left and right insets, arrows). Scale bar for full tomograms 200 nm, scale bar for insets 25 nm.

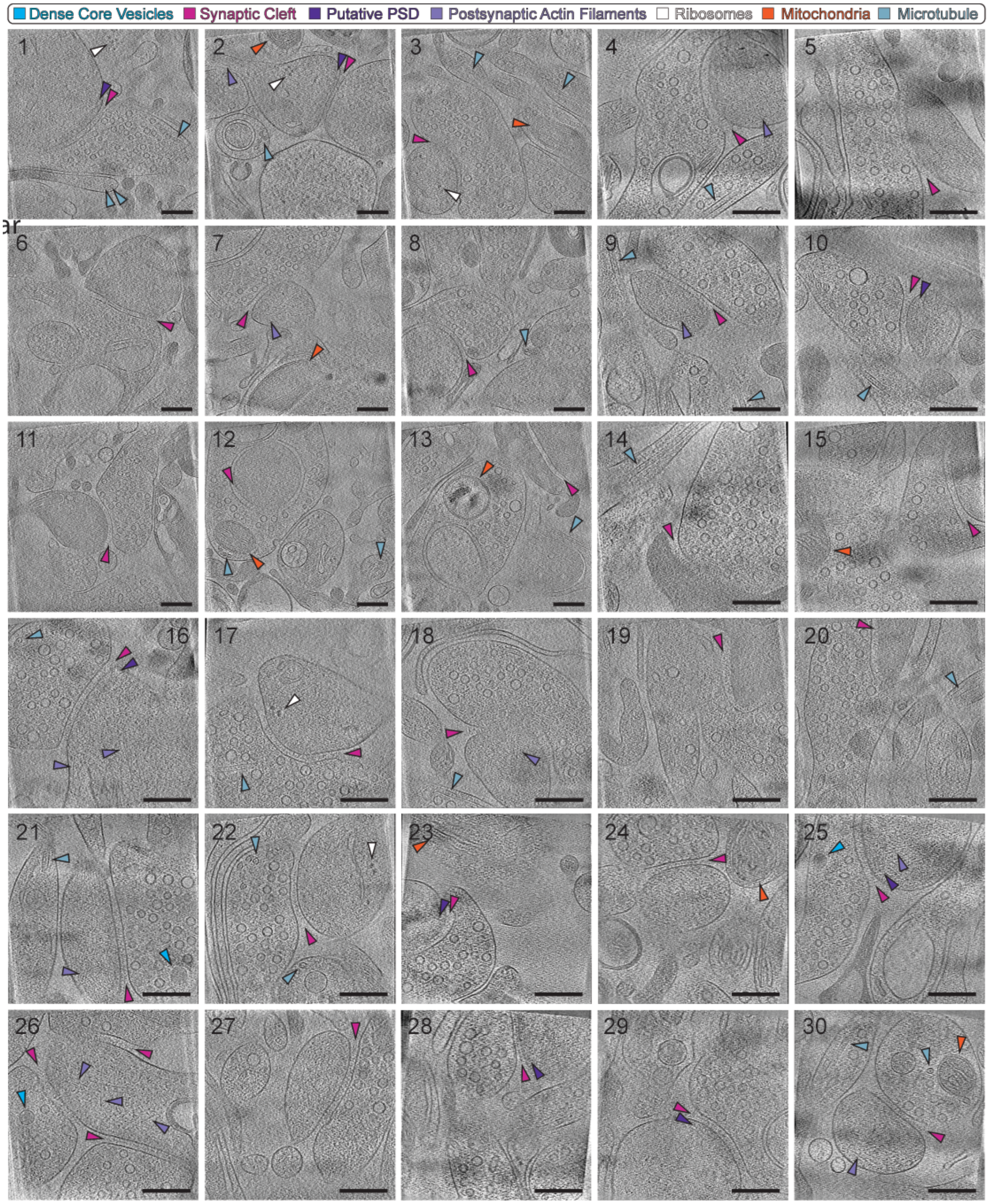


**Extended Data Figure 7. Synapse diversity in the CA1-sr.** An array of 30 out of 107 total synapses representing all datasets including synapses from the planar lift-out (23-25, 28-30). Arrows highlight the synaptic cleft (pink) in all tomogram cross sections, dense core vesicles (blue) in the presynaptic terminal, putative postsynaptic density (PSD, dark purple) when present, postsynaptic actin filaments (light purple), ribosomes (white), mitochondria (orange) and microtubules (teal). All scale bars 200 nm.

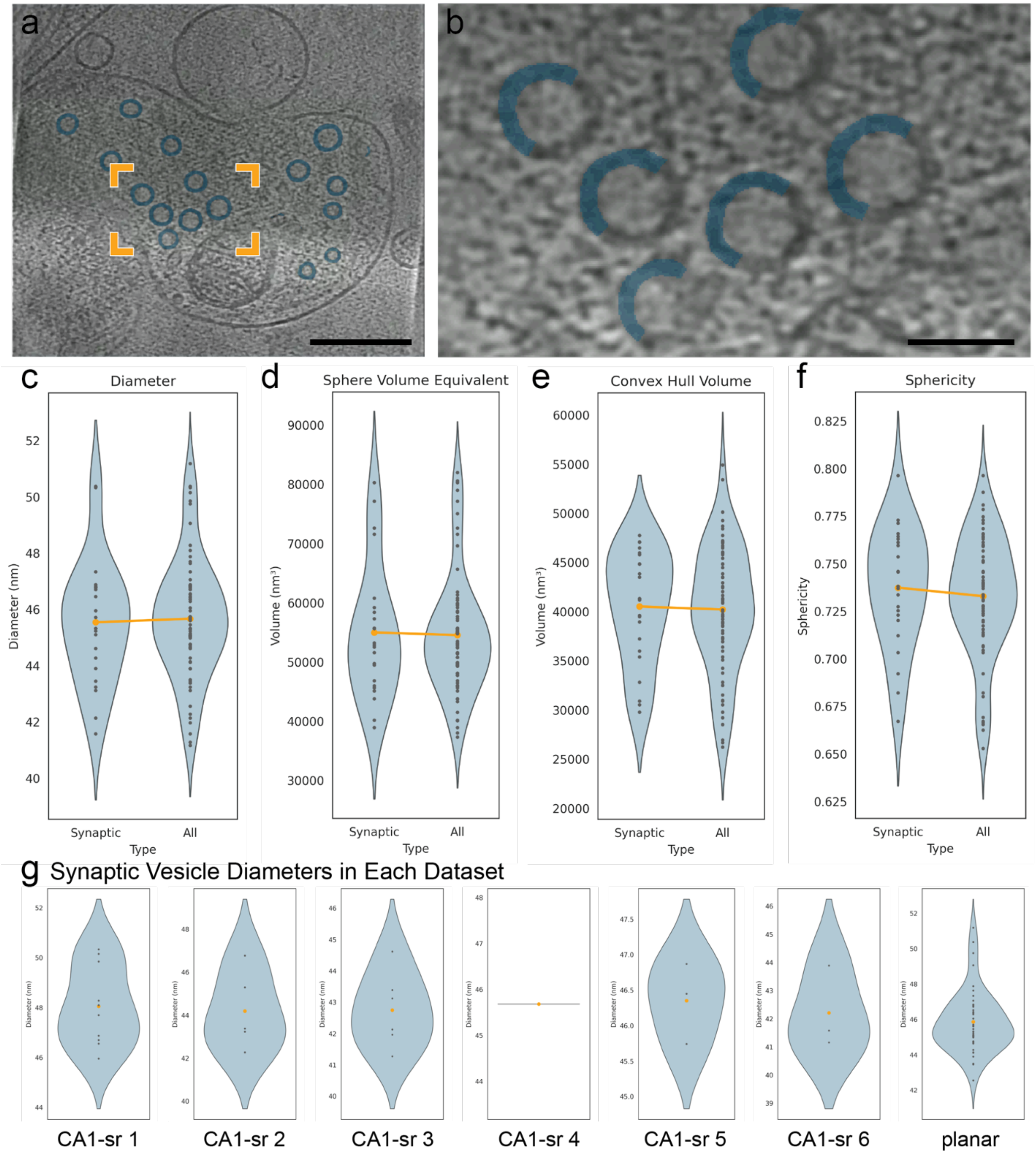


**Extended Data Figure 8. Segmentation and Size Distribution of Synaptic Vesicles**. a) The image shows a tomogram with 16 visible segmentations (62 vesicles in total throughout the tomogram), scale bar 200 nm. b) A crop of the tomogram with 6 vesicles, where half of the vesicle has been removed to show the degree to which the segmentation lines up with the vesicle boundaries, scale bar 50 nm. c) Violin plot of the mean diameter, with left plot (Synaptic) including only tomograms where all vesicles are in a visible synapse and the right plot (All) includes all vesicles in all synapse tomograms. The diameter is measured in 2d, per slice, where dots are the per-tomogram means of the median of the per-slice 2d feret diameter of each vesicle. The median was chosen to be more robust to outliers.

d) Plot of the mean Sphere Volume Equivalent per tomogram, which is the calculated volume of a sphere, given the diameter used in (c). e) The mean volume of the convex hull of the segmented connected component, per tomogram. The segmented component is a is the shell of the vesicle, so the convex hull is used to fill in the interior. f) A violin plot of the mean sphericity of the vesicles, per tomogram. g) Synaptic vesicle diameter distribution for each dataset.

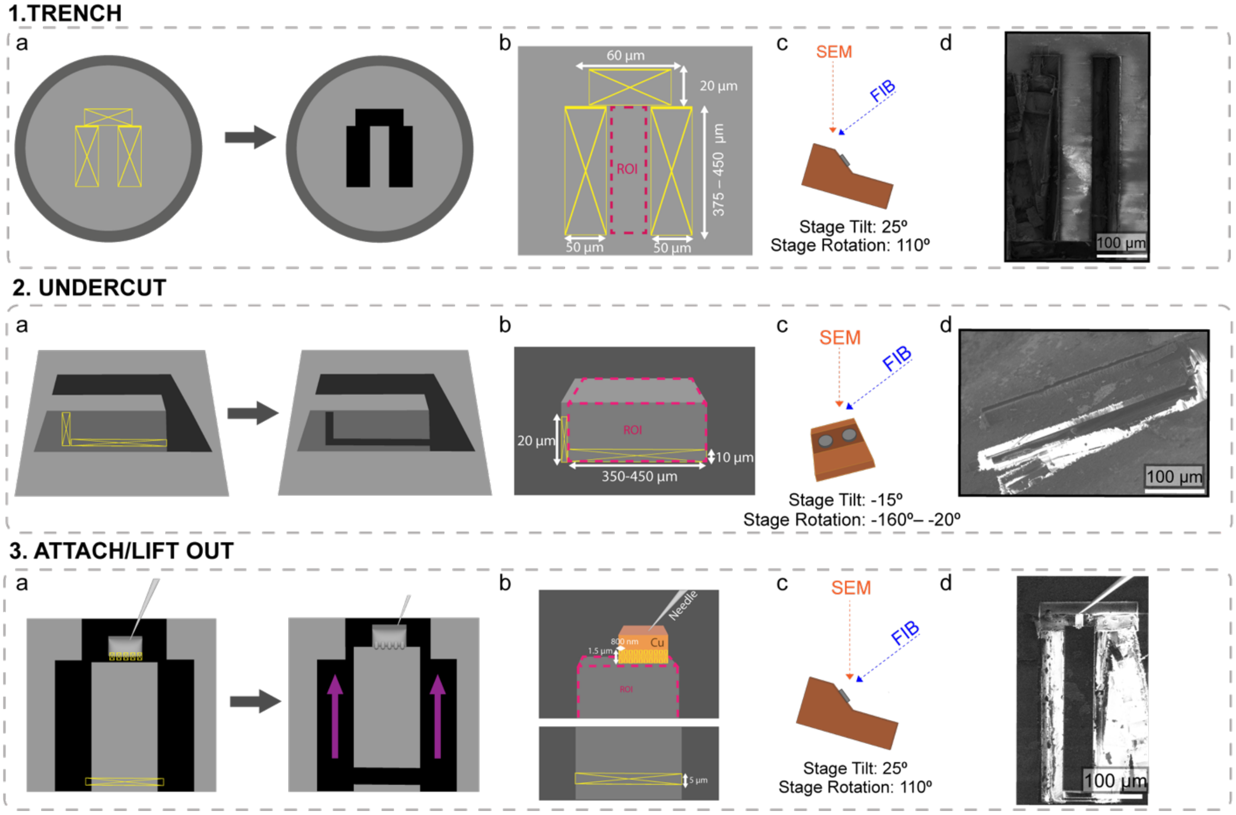


**Extended Data Figure 9. Milling Procedure for Cryo-Lift Out of Planar Samples from HPF Carriers.** 1a, 2a & 3a) Schematic diagram showing the FIB milling patterns (yellow) performed on the sample for Trench Milling, Undercutting and Attachment & Lift Out steps respectively. For the Attachment & Lift Out step, the lift out needle is depicted on top of the region of interest and the purple arrow denotes the direction of the lift out. 1b, 2b & 2c) The region of interest (ROI) (Pink) and the dimensions of FIB milling patterns (yellow) placed for Trench Milling, Undercutting and Attachment & Lift Out steps respectively. 1c, 2c & 3c) Schematic diagram for the orientation of the SEM and FIB beams in relation to the FIB sample holder, with stage tilts and rotation angles for Trench Milling, Undercutting and Attachment & Lift Out steps respectively. 1d, 2d and 3d) Examples of FIB images for Trench Milling, Undercutting and Attachment & Lift Out steps respectively

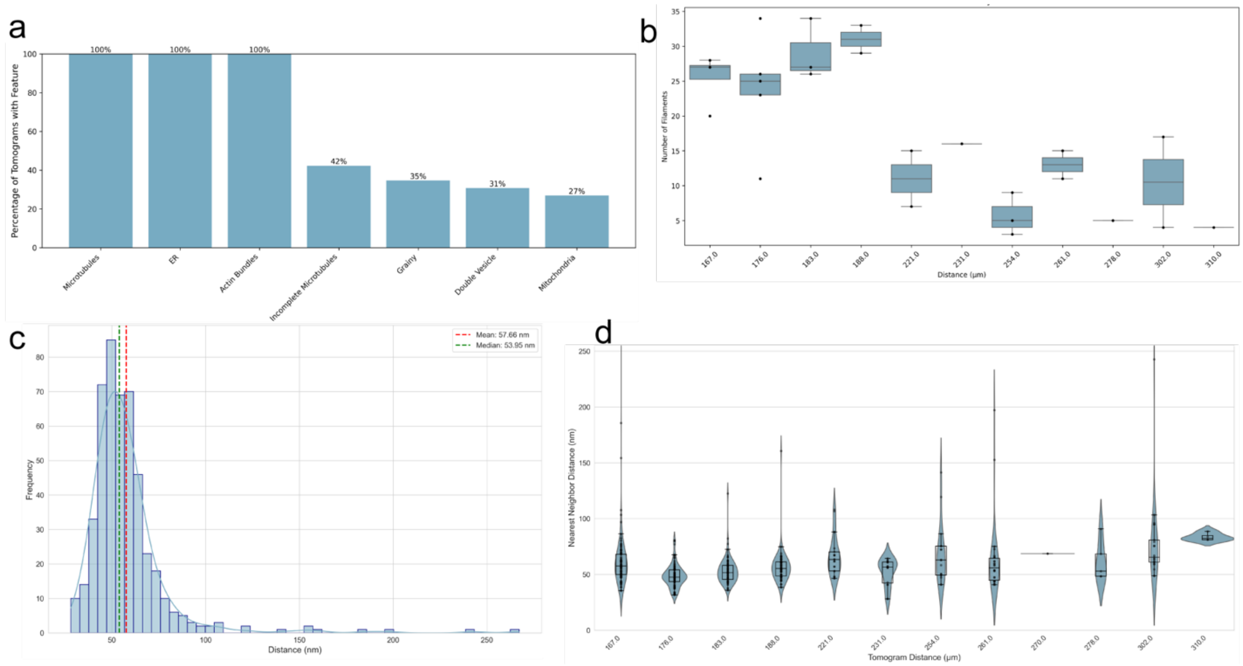


**Extended Data Figure 10. Analysis of the cytoskeletal composition and organisation of the apical dendrite network.** a) Characterisation of the percentage of the cellular components observed in 26 reconstructed tomograms depicting axial views of dendrites. b) The number of microtubules observed in tomograms as a function of distance from the CA1so. c) Histogram of the nearest neighbour differences between microtubules from all of the 26 dendrite tomograms. Mean and median values are plotted as red and green dashed lines respectively. d) Violin plot of nearest neighbour distances between microtubules as a function of distance from the CA1so.

| Mouse Age (days) | Brain Region | Section Thickness (µm) | Carrier Material | Carrier Recess Depth (µm) | Cryo Protectant and Buffer | Incubation Time (min) | Vitreous Rating | E.D. Figure 3 Number |
|---|---|---|---|---|---|---|---|---|
| 14 | Cortex | 200 | Aluminum | 200 | 20% Dextran+ in PBS pH 7.4 | 0 | X | 1 |
| 13 | Cortex | 150 | Aluminum | 150 | 20% Dextran in PBS pH 7.4 | 30 | X | 2 |
| 7 | Cortex | 150 | Copper | 200 | 20% Dextran in PBS pH 7.4 | 30 | X | 3 |
| 118 | Cortex | 150 | Copper | 200 | 20% BSA in 100 mM PB pH 7.4 | 35 | X | 4 |
| 118 | Cortex | 100 | Copper | 100 | 20% BSA in 100 mM PB pH 7.4 | 0 | X | 5 |
| 9 | Cortex | 150 | Copper | 200 | 10% Dextran 5% Sucrose in 100 mM PB pH 7.4 | 25 | X | |
| 38 | Cortex | 100 | Copper | 100 | 10% Dextran 5% Sucrose in 100 mM PB pH 7.4 | 20 | +* | |
| 125 | Cortex | 100 | Copper | 100 | 10% Dextran 5% Sucrose in 100 mM PB pH 7.4 | 25 | ± | 6 |
| 125 | Cortex | 100 | Copper | 100 | 10% Dextran 5% Sucrose in 100 mM PB pH 7.4 | 30 | ± | 7 |
| 125 | Cortex | 100 | Copper | 100 | 10% Dextran 5% Sucrose 5% Ethylene Glycol in 100 mM PB pH 7.4 | 20 | + | 8 |
| 149 | Cortex | 100 | Copper | 100 | 10% Dextran 5% Sucrose 5% Ethylene Glycol in 100 mM PB pH 7.4 | 25 | ++ | 9 |
| 125 | Cortex | 100 | Copper | 100 | 10% Dextran 10% Sucrose in 100 mM PB pH 7.4 | 20 | ++ | 10 |
| 149 | Cortex | 100 | Copper | 100 | 10% Dextran 10% Sucrose in 100 mM PB pH 7.4 | 25 | ++* | 11 |
| 184 | Hippocampus | 100 | Copper | 100 | 10% Dextran 10% Sucrose in NMDG pH 7.4 | 0 | X | 12 |
| 184 | Hippocampus | 100 | Copper | 100 | 10% Dextran 10% Sucrose in NMDG pH 7.4 | 15 | ++ | 13 |
| 184 | Hippocampus | 100 | Copper | 100 | 10% Dextran 5% Sucrose 5% Ethylene Glycol in aCSF[a] pH 7.4 | 20 | ++ | 14 |
| 144 | Cortex | 100 | Copper | 100 | 20% Dextran in NMDG pH 7.4 | 25 | X | 15 |
| 144 | Cortex | 200 | Copper | 200 | 10% Dextran 10% Sucrose in NMDG pH 7.4 | 30 | ++ | 16 |

[+]Dextran 40,000 MW

*Conclusion based on one tilt series

[a]aCSF composition 119 mM NaCl, 26.2 mM NaHCO3, 2.5 mM KCl, 1 mM Na2HPO4, 1.3 mM MgCl2, 10 mM glucose, 2.5 mM CaCl2

**Extended Data Table 1. Conditions assessed for vitrification state.**

| Step | Sub-step | Current | Pattern Type | Pattern Dimensions (µm) | | | Time (min) | Notes |
|---|---|---|---|---|---|---|---|---|
| | | | | X | Y | Z | | |
| Fluorescence Overview | NA | NA | NA | NA | NA | NA | 20-50 | Time for imaging with 1 fluorophore, 20 minutes for 3 Z steps, 50 minutes for 12 Z steps. |
| Fiducials | NA | 4-60 nA | Rectangle | 75 | 4 | 3 | 10 | Used for alignment between SEM and IFM. |
| Trenches: Perpendicular | Long Trench | 60 nA | RCS | 60 | 150-200 | 5-6 | <45 | For a desired lift-out length of ~60-70 um. Longest long trenches should be used in conjunction with deepest deep trenches. For 100 um thick samples, long trench lengths <170 um and deep trench depths <9 should be used to avoid milling into the metal carrier. Largest lift-outs yield ~10-14 3-5 µm thick sections. RCS patterns mill towards target region. |
| | Deep Trench | 60 nA | RCS | 60 | 40-60 | 6-11 | | |
| Trenches: Planar | Side Trenches | 200 nA | RCS | 50 | 350-400 | 6 | 7–8 | For a desired lift-out length of ~350-400 µm. Trenches are milled symmetrically at either side of the region of interest, with the scan pattern directed towards the area for lift out. |
| | Top Trench | 60 nA | RCS | 210 | 50 | 2 | 6 | To allow the lift-out needle with the copper block attachment to be brought to the top of the sample. The scan pattern is directed towards the area for lift out. |
| Side and Undercuts: Perpendicular | Side Cuts | 15 nA | Rectangle | 4 | 40-60 | 4 | 10 | Use a stage tilt of <13º to avoid milling into the metal carrier for deep lift-outs. |
| | Undercut | 15 nA | Rectangle | 53-58 | 6-8 | 4 | | |
| Side and Undercuts: Planar | Undercut | 15 nA | Rectangle | 350 - 400 | 6 | 1 | 16–20 | Perform undercuts from both ±90º stage rotation relative to the region of interest. The time listed is for milling from one of the angles. |
| Weld Copper Block to Tissue | NA | 0.3 nA | CCS | 1 | 2 | 3* | < 5 | Tilt the stage to the shallowest angle where remaining material from side cuts is still visible, roughly -3 to 3º . Milling patterns should be oriented to mill from tissue towards the copper block. After attachment, then mill away remaining side material. Dwell time was set to 30 µs |
| Sectioning and Welding | Sectioning | 1-4 nA | Line | ~50 | NA | 4 | 2-5 | The stage was tilted to the shallowest angle possible, around –3 - -5º, for section deposition. 4 nA allows for faster cuts but 1 nA results in a smoother surface that is advantageous for subsequent thinning steps. |
| | Welding | 0.3 nA | CCS | 3 | 0.8 | 5 | 3-4 / section | Stage was tiled to 15º before placing welding patterns. Welds were placed with 4 um periodicity such that each section had 3-4 welds per side with 6-8 welds per section in total. CCS patterns milled from the section towards the copper bars. Dwell time was set to 30 µs and each welding pattern took 30 seconds to mill. |

(*) Using a milling application based on the Xe sputter yield

**Extended Data Table 2 – Trench milling parameters for perpendicular lift-outs.**

| Mouse Sex | Mouse Age | Hemisphere (L/R) | Cryo Protectant and Buffer | Incubation Time (min) | Region | Dataset Number (for CA1-sr) | Tilt Series Collected | Synapses | % Dataset that is Synapses | Average Synaptic Vesicle Diameter (nm) |
|---|---|---|---|---|---|---|---|---|---|---|
| M | 184 | Not recorded | 10% Dextran, 10% Sucrose in NMDG, pH 7.4 | 15 | CA1-sr | 1 | 58 | 16 | 28 | 48.0 |
| M | 184 | L | 10% Dextran, 5% Sucrose, 5% Ethylene Glycol in aCSF, pH 7.4 | 20 | CA1-sr | 6 | 10 | 3 | 30 | 42.2 |
| M | 184 | L | 10% Dextran, 10% Sucrose in aCSF, pH 7.4 | 20 | CA1-sr | 5 | 22 | 5 | 23 | 46.4 |
| F | 172 | R | 10% Dextran, 10% sucrose in NMDG, pH 7.4 | 20 | CA1-sr | 2 | 145 | 19 | 13 | 44.1 |
| M | 184 | R | 10% Dextran, 10% sucrose in NMDG, pH 7.4 | 15 | CA1-sr | 3 | 60 | 11 | 18 | 42.7 |
| F | 172 | R | 10% Dextran, 10% sucrose in NMDG, pH 7.4 | 20 | CA1-sp | | 35 | 0 | 0 | N/A |
| F | 172 | R | 10% Dextran, 10% sucrose in NMDG, pH 7.4 | 20 | CA1-sr | 4 | 64 | 2 | 3 | 45.7 |
| M | 184 | L | 10% Dextran, 10% sucrose in NMDG, pH 7.4 | 15 | CA1* | | 252 | 51 | 20 | 45.9 |
| F | 144 | Not recorded | 10% Dextran, 10% sucrose in NMDG, pH 7.4 | 30 | cortex | | 28 | 0 | 0 | N/A |

**Extended Data Table 3. Datasets Collected.** Table of all vitreous datasets collected in this work.
